# Microbiome depletion rejuvenates the aging brain

**DOI:** 10.64898/2026.02.13.705770

**Authors:** C. Gasperini, K.M. Holton, F. Limone, M. Juttu, C.C. DeMeo, K. Kekrtova, S. Patankar, K.M. Wells, R.M. Giadone, L. Ben Driss, G. Wei, A. Kiem, Q. Xu, R.T. Lee, M. Friedlander, D.T. Scadden, L.L. Rubin

**Affiliations:** Department of Stem Cell and Regenerative Biology, Harvard University, Cambridge, MA 02138, USA; Harvard Stem Cell Institute, Cambridge, MA 02138, USA; Broad Institute of MIT and Harvard, Cambridge MA 02142, USA; Institute for Translational Neuroscience, NYU Grossman School of Medicine, New York, NY, USA; National Institute of Mental Health, Topolova 748, Klecany 250 67, Czech Republic; Charles University, Third Faculty of Medicine, Ruska 87, Prague 100 00, Czech Republic; Department of Molecular Medicine, The Scripps Research Institute, La Jolla, CA, 92037, USA; Lowy Medical Research Institute, La Jolla, California 92037, USA; Center for Regenerative Medicine, Massachusetts General Hospital, Boston, MA, 02118, USA; Department of Biomedical Informatics, Harvard Medical School, Boston, MA, 02115, USA

## Abstract

Aging is associated with cognitive decline and increased vulnerability to neurodegeneration driven by an array of molecular and cellular changes like impaired vascular integrity, demyelination, reduced neurogenesis, and chronic inflammation. Recent studies implicate the gut microbiome as a modulator of brain aging, but the underlying mechanisms remain elusive. Here, we show that depleting the gut microbiome by administering antibiotics to aged mice induces widespread molecular and structural rejuvenation in the brain. Our transcriptomic analyses by single-nucleus RNA sequencing revealed pronounced transcriptional shifts across multiple brain cell types. We confirmed that antibiotic treatment improves vascular density, promotes myelination, enhances neurogenesis, and reduces microglial reactivity. Functionally, microbiome-depleted mice showed improved hippocampal memory performance. Analyses of brain and plasma cytokine levels showed a decrease in several pro-inflammatory factors post-treatment and identified candidate factors, including the chemokine eotaxin-1. Inhibiting eotaxin-1 alone can reverse several aspects of brain aging. Our findings demonstrate that age-associated microbial inflammation contributes to brain aging and that its attenuation can restore youthful features at the molecular, cellular, and functional levels. Targeting the gut microbiome or its circulating mediators may therefore represent a non-invasive approach to promote brain health and cognitive resilience in aging.

## Introduction

The global population aged 60 years and older is projected to increase from about 12% in 2015 to approximately 22% by 2050 ^1^. Despite a substantial increase in life expectancy, people are living longer, but not necessarily healthier lives, spending an increasing number of years in poor health. Aging is the greatest risk factor for many of the most prevalent human diseases, including cancer, diabetes, and, especially, neurodegenerative disorders ^2^. The incidence of Alzheimer’s disease (AD), Parkinson’s disease (PD), and amyotrophic lateral sclerosis (ALS) rises dramatically with age, presenting growing challenges to public health and healthcare systems ^2^.

Understanding the biological mechanisms that drive aging and neurodegeneration is therefore a pressing priority. Aging is characterized by a progressive accumulation of molecular and cellular damage, reflected in conserved changes, commonly referred to as hallmarks, such as genomic instability, epigenetic alterations, mitochondrial dysfunction, cellular senescence, chronic inflammation, loss of proteostatic control, and dysbiosis that occur in many cell types ^3,4^, along with cell-specific changes ^5^. In the central nervous system (CNS), reduced synaptic plasticity and neurogenesis have been observed, while oligodendrocyte dysfunction impairs myelin maintenance and axonal integrity ^6^. Age-related vascular decline is associated with reduced vessel density and a compromised blood brain barrier, further contributing to a reduction in brain health. These degenerative processes are accompanied by increased neuroinflammation, evidenced by an the appearance of reactive microglia, gliosis, and elevated pro-inflammatory cytokine levels ^7^. These changes impair tissue homeostasis and resilience, while negatively affecting a variety of behaviors.

Although traditionally viewed as irreversible, aging is increasingly recognized as a malleable process. Over the past four decades, experimental models have demonstrated that systemic manipulations can partially restore youthful function in aged tissues ^8^. One of the most compelling examples is heterochronic parabiosis, in which the circulatory system of a young and an old mouse are surgically joined. This model has shown that exposure to young blood can rejuvenate aged tissues, including the brain^9–12^. In particular, our group and others have demonstrated that heterochronic parabiosis improves neurogenesis, vascular integrity, and even behavioral performance in aged mice ^10,12^. Recent single-cell analyses have further revealed that young blood exposure reprograms aging-related transcriptional signatures in most cell types, culminating in a reduction in senescence of multiple cell types ^10^. Beyond parabiosis, other systemic interventions such as plasma transfer and exercise restore regenerative capacity and cognitive performance in aged mice ^13–15^. Perhaps surprisingly, these complex manipulations have led to the identification of a variety of circulating factors that, when administered systemically, positively affect the aged CNS.

Emerging evidence suggests that the gut microbiome, a densely populated microbial ecosystem containing up to 100 trillion organisms, may play a central role in CNS aging. Microbiome composition undergoes dynamic changes across the lifespan. It develops alongside the immune and nervous systems in early life, stabilizes in adulthood, and is disrupted in aging. Age-related dysbiosis is marked by reduced microbial diversity, expansion of pro-inflammatory taxa, and impaired intestinal barrier integrity, which together lead to increased levels of circulating microbial metabolites and systemic inflammation. This chronic inflammatory state has been linked to frailty, metabolic dysfunction, and neurological disease ^16^. Dysbiosis is also a consistent feature in neurodegenerative conditions such as AD, PD, and ALS ^17^, and microbiome-targeting interventions have shown modest benefit in preclinical models ^18,19^. In those cases, how the changing gut microbiome ultimately affects the brain is incompletely understood.

Here, we test the hypothesis that targeting the age-associated gut microbiome can counteract key features of brain aging. To do so, we used oral antibiotics to deplete the microbiota of aged mice and performed a comprehensive, multi-level analysis of brain and systemic phenotypes. Remarkably, microbiome depletion induced widespread rejuvenation across cortical regions and multiple brain cell types, including improved cerebrovascular integrity, characterized by increased vessel density and enhanced expression of tight junction components, as well as elevated oligodendrocyte maturation and myelin production. In parallel, we observed increased neural stem cell proliferation and enhanced neurogenesis in the hippocampus. At the molecular level, single-nucleus RNA sequencing revealed transcriptional reprogramming consistent with a reversal of aging hallmarks across diverse neural and non-neural cell types. These brain-specific effects are accompanied by systemic changes, including reduced inflammatory signaling in the brain and in peripheral organs and decreased circulating levels of pro-aging factors. Among these, the chemokine eotaxin-1 (CCL11) emerged as a critical effector: its inhibition is sufficient to mimic the beneficial effects of microbiome depletion on the aging brain, while its overexpression blunts them.

Together, these findings identify the aging gut microbiome as a modifiable regulator of brain and systemic aging. By restoring microbial balance, it is possible to reactivate regenerative and homeostatic programs, highlighting the gut–brain axis as a promising therapeutic target to promote healthy aging and cognitive resilience.

## Results

### Microbiome depletion induces global transcriptional shifts in the aged brain

To determine whether targeting the gut microbiome can modulate brain aging, we depleted the gut microbiota of 24-month-old mice using a well-established cocktail of four antibiotics (Abx: ampicillin, neomycin, vancomycin and metronidazole) ^17,20^. These antibiotics were selected for their broad-spectrum activity, collectively targeting Gram-positive and Gram-negative bacteria as well as anaerobes, thereby ensuring effective and sustained depletion of the gut microbiota ^21^.

Antibiotics were administered via oral gavage twice daily for 7 days to ensure direct delivery to the gut, followed by administration in drinking water for an additional 21 days to maintain depletion (Fig. 1A). Metronidazole was omitted during the drinking water phase to prevent potential confounding effects, as this compound is known to cross the blood–brain barrier ^22^. Control mice received water following an identical schedule. 16S rRNA gene sequencing of bacterial diversity in fecal samples confirmed robust depletion of the gut microbiota. A marked reduction in alpha diversity, reflecting loss of taxonomic richness and evenness, was observed (Extended Data Fig. 1A). Moreover, operational taxonomic unit (OTU) analysis revealed a dramatic decline in the abundance of closely related bacterial groups in Abx-treated mice (Fig. 1B and Extended data Fig 1B), indicating efficient microbiome disruption.

**Fig 1.**
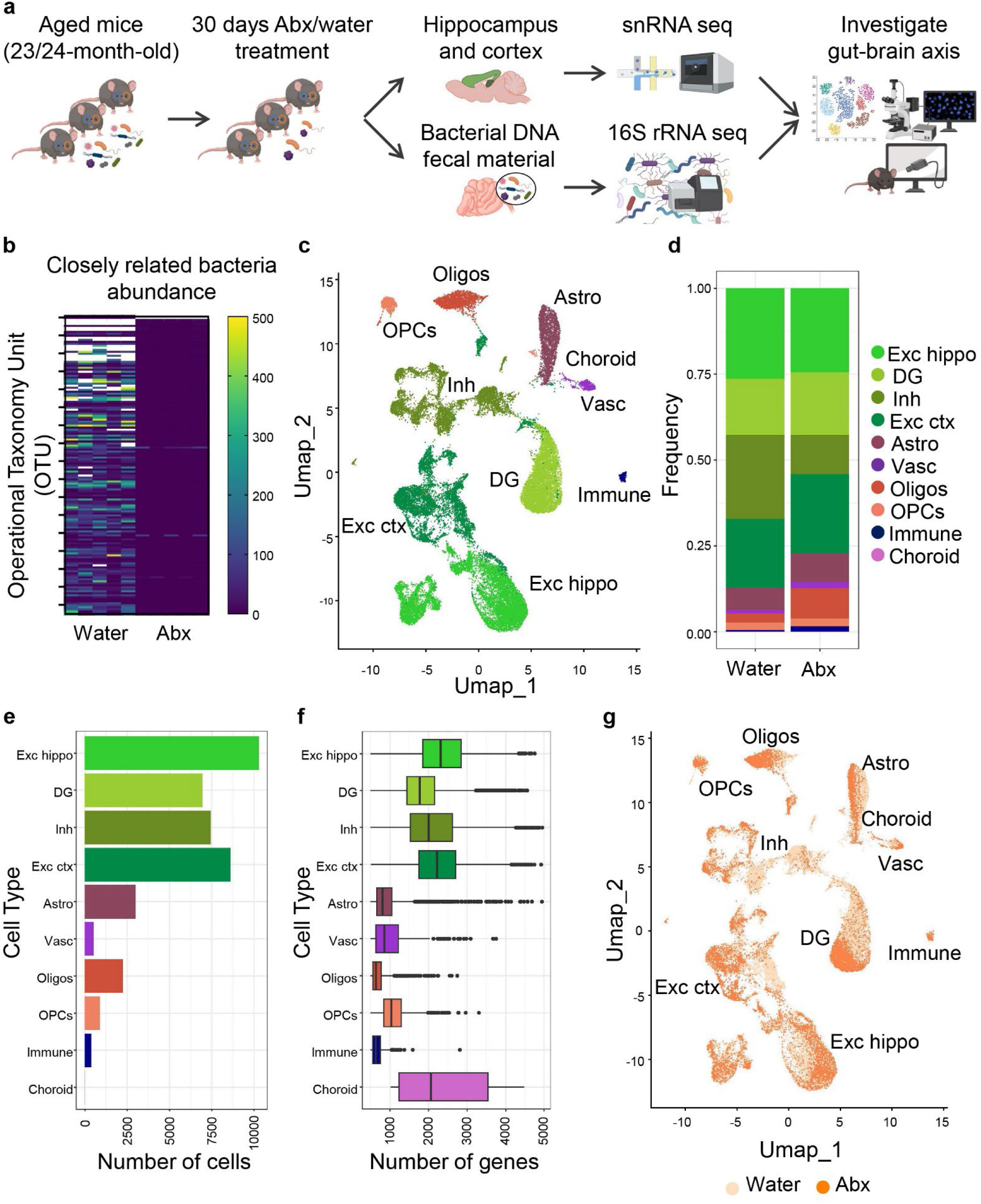
Microbiome depletion triggers shift in brain cells transcriptional profile. **a**, Schematic of experimental design: 24-month-old mice were treated with a cocktail of four antibiotics (Abx) for 30 days to deplete the gut microbiota. Brains, fecal samples, and peripheral tissues were collected for molecular analyses. **b**, Heatmap showing operational taxonomic unit (OTU) abundance in fecal samples from water- and Abx-treated mice, indicating a marked reduction in bacterial diversity and richness. **c**, UMAP embedding of 45,089 single-nucleus transcriptomes from cortex and hippocampus, identifying ten major brain cell types based on marker gene expression. **d**, Relative frequency of each cell type in water-and Abx-treated groups shows no major shifts in cell-type composition. **e**, Total number of cells detected per cluster, revealing neuronal enrichment consistent with dissected regions. **f**, Distribution of detected genes per nucleus across cell types; excitatory hippocampal and cortical neurons, inhibitory neurons, and DG granule cells show high transcriptional complexity. **g**, UMAP embedding split by treatment group shows that microbiome depletion alters transcriptional signatures across multiple brain cell types.

Mice were monitored daily, and no significant changes in body weight were observed during treatment (Extended Data Fig. 1C). After 30 days, mice were perfused with PBS, and multiple organs were harvested for analysis. In line with previous rejuvenation paradigms ^23–25^, we observed reduced relative weights of the liver, heart, and white adipose tissue, as well as the characteristic enlargement of the cecum following microbiome depletion, reflecting unprocessed dietary fiber accumulation (Extended Data Fig. 1D-F).

To assess the consequences of microbiome depletion on the aging brain, we performed highthroughput single-nucleus RNA sequencing on hippocampal and cortical samples from Abx- and water-treated mice. We choose these two regions because the hippocampus is a neurogenic niche where adult neurogenesis markedly declines with age ^26^, and the cortex contains a broad diversity of brain cell types, therefore enabling comprehensive cellular profiling ^27^. Brain tissues were processed using a previously validated protocol ^28^ yielding transcriptomic profiles from 45,089 nuclei after quality control. Using canonical marker genes ^29^, we identified ten major brain cell types, including excitatory neurons (Exc) from the cortex (Ctx), hippocampus (HC) and dentate gyrus (DG), inhibitory (Inh) neurons, oligodendrocytes (oligos), oligodendroctye precursor cells (OPCs), astrocytes (astro), immune cells (immune), vascular cells (Vasc), and choroid plexus epithelial cells (choroid) (Fig. 1C and Extended Data Fig. 2). Marker gene analysis confirmed the identity of each cluster, and unsupervised clustering revealed representation of all major populations across both treatment groups (Fig. 1D). Quantification via propeller showed that the relative abundance of cell types remained stable between conditions at an FDR of 0.05 (Fig. 1D), with the predominance of neurons due to the hippocampal and cortical origin of the samples (Fig. 1E). Among all populations, excitatory cortical and hippocampal neurons, inhibitory neurons, and dentate gyrus granule cells exhibited the highest numbers of detected genes per cell, indicative of transcriptional complexity (Fig. 1F). Although choroid plexus epithelial cells expressed over 2,000 genes, they were not consistently represented across individual mice and were excluded from downstream analyses. Microbiome depletion induced marked shifts in the transcriptional profiles of multiple brain cell types, suggesting widespread modulation of gene expression in the brain (Fig. 1G). These findings prompted further high-resolution subclustering analyses to dissect the impact of microbiome depletion on specific cell states and brain functions.

### Microbiome depletion improves vasculature integrity in the aging brain

Given the widespread transcriptional reprogramming observed across multiple brain cell types following microbiome depletion (Fig. 1), we next sought to investigate which specific biological processes were most affected. We started by focusing on brain vasculature, a compartment highly susceptible to aging, intimately involved in neuroimmune and neurovascular signaling and the key interface between blood and brain ^30^, for deeper transcriptional and structural characterization. Subclustering of vascular cells from single-nucleus RNA-seq data revealed that microbiome depletion markedly reprogrammed their transcriptional profiles, resulting in a distinct redistribution of cell states (Fig. 2A). Functional enrichment analysis of differentially expressed genes (DEGs) pointed to significant changes in vascular-related pathways, including endothelial cell migration, angiogenesis, morphogenesis, and blood-brain barrier function (Fig. 2B). Among the most differentially expressed genes were pro-angiogenic and vascular remodeling factors such as Clic4, Adgrl4, and Plxna2, as well as other genes implicated in endothelial barrier function, including Slc2a (Fig. 2C). Conversely, expression of several anti-angiogenic and vascular-destabilizing factors (Clu, Ptn, and Cst3), was significantly reduced in microbiome-depleted animals, suggesting a shift toward a more pro-angiogenic transcriptional state (Fig. 2C).

**Fig 2.**
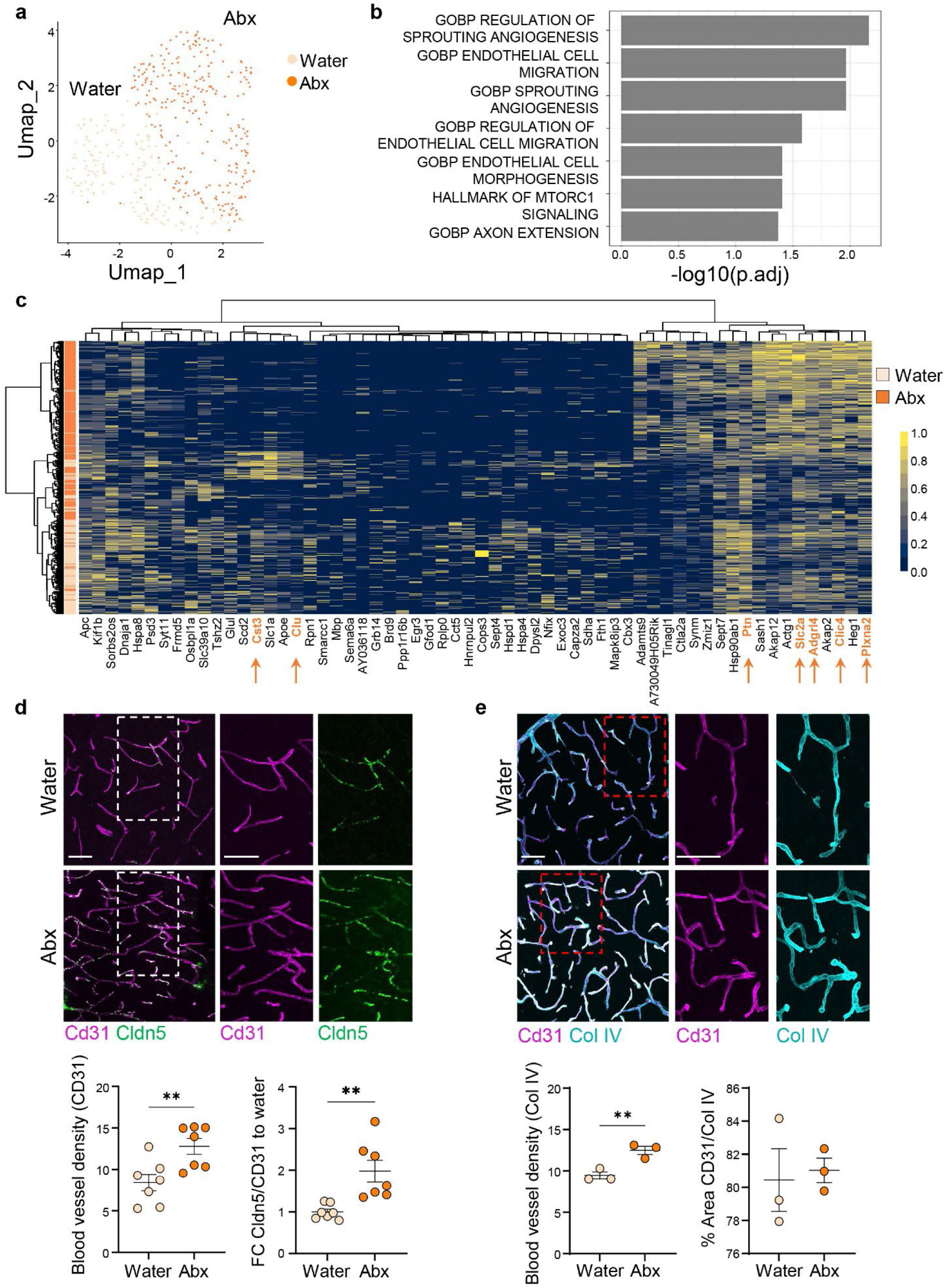
Microbiome depletion triggers changes in brain vasculature. **a,** UMAP embedding of endothelial cells colored by treatment group shows a distinct transcriptional shift upon microbiome depletion. **b,** Over representation analysis of differentially expressed genes reveals enrichment in pathways related to angiogenesis, endothelial migration, and vascular morphogenesis. **c,** Heatmap of differentially expressed genes in endothelial cells between Abx- and water-treated mice. **d,** Representative immunofluorescence images of CD31+ blood vessels (magenta) and tight junction marker Claudin-5 (green) in cortical sections; bottom, quantification of vessel density and CD31+ Cldn5+ /CD31+ area (n = 7/group). **e,** Representative low-magnification images of cortical vasculature with high-magnification insets (dashed boxes) showing CD31(magenta) and Collagen IV (cyan) staining; bottom panel, quantification of blood vessel density as Collagen IV + area, and percentage of colocalization Cd31+ Col IV+ area (n = 3/group). Scale bar, 50 μm. All data represented as mean ± s.e.m.; \*\**P* < 0.01, Student’s *t*-test.

To determine whether these transcriptional changes translated into structural alterations in the cerebral vasculature, we performed immunofluorescence analysis using established endothelial and tight junction markers. Cortical sections stained for CD31, which labels endothelial cells, and Collagen IV, a major component of the vascular basement membrane, showed a significant increase in blood vessel density in Abx-treated mice compared to controls (Fig. 2D, E). Furthermore, expression of Claudin-5 (Cldn5), a key tight junction component of the blood–brain barrier, was markedly elevated in the Abx group, indicating enhanced vascular integrity (Fig. 2D). Interestingly, while the percentage Collagen IV⁺ endothelial cell segments that was also CD31^+^ was not significantly different between groups (Fig. 2E), water-treated mice displayed rare CD31⁻/Collagen IV⁺ profiles (inset), consistent with residual basement membrane structures lacking endothelial coverage, a feature commonly associated with vascular aging. In contrast, Abx-treated mice showed more uniform CD31⁺/Collagen IV⁺ staining (Fig. 2E, inset). These changes were not due to significant differences in overall cell number, as indicated by the sequencing data (Fig. 1D), supporting a direct effect on vascular remodeling.

Together, these findings indicate that microbiome depletion promotes vascular remodeling and strengthens the endothelial barrier in the aging brain, potentially mitigating vascular deterioration associated with age and inflammation.

### Microbiome depletion enhances myelin maintenance in old brains

Another hallmark of brain aging is the progressive loss of myelin across multiple brain regions, leading to impaired signal transduction and synaptic dysfunction ^3^. To assess whether microbiome depletion affects oligodendrocyte transcriptional states, we performed subclustering and dimensionality reduction using UMAP. Abx treatment induced a clear shift in the distribution of cells within these clusters, suggesting that microbiome depletion modulates gene expression in oligodendrocyte lineage cells (Fig. 3A). Functional enrichment analysis of DEGs identified significant terms involved in neuron projection development, axonal extension, and synaptic structure and function (Fig. 3B). Consistently, DEG analysis revealed that microbiome depletion upregulated transcripts associated with oligodendrocyte differentiation and maturation (Cdh8, Daam2, Lama2) as well as myelin formation (Gli2, Dlg2, Il33), while reducing negative regulators of myelination (Apoe, Epha4) (Fig. 3C, D).

**Fig 3.**
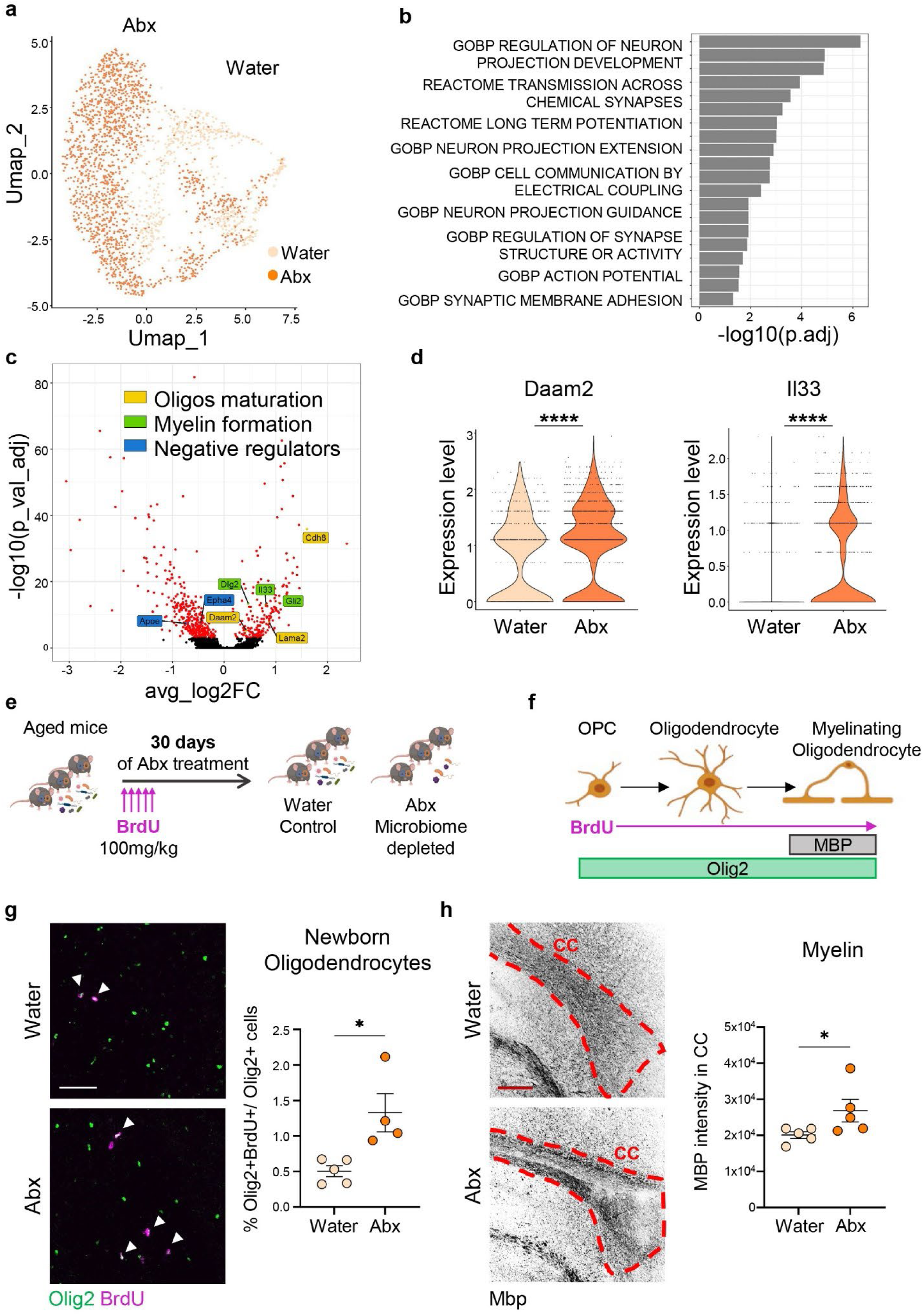
Microbiome depletion triggers myelin production in old brains. **a,** UMAP plot showing re-clustered oligodendrocytes colored by treatment group; Abx treatment induces a shift in transcriptional state without generating new subtypes. **b,** Over representation analysis of differentially expressed genes highlights pathways related to neuronal projection development, synaptic signaling, and axonal structure. **c,** Volcano plot of differentially expressed genes in oligodendrocytes between Abx- and water-treated mice. X-axis is the log2 foldchange, Y-axis is the log10 of the Bonferonni adjusted p-value. **d,** Violin plots showing increased expression of Daam2 and Il33, genes involved in oligodendrocyte maturation and myelination, in Abx-treated mice. **e,** Experimental design for BrdU incorporation during microbiome depletion to label dividing progenitors. **f,** Schematic of oligodendrocyte maturation from OPCs, and marker expression used to track differentiation and myelination (BrdU, Olig2, MBP). **g,** Representative images and quantification of BrdU+ (magenta) Olig2+ (green) cells in the corpus callosum, indicating increased generation of newborn oligodendrocytes in Abx-treated mice. White arrows indicate double positive cells. **h,** Representative images and quantification of MBP intensity in the corpus callosum, highlighted in red, showing enhanced myelin content following microbiome depletion (n = 5/group). Scale bar, 50 μm (g) and 100 μm (h). All data represented as mean ± s.e.m.; \**P* < 0.05, **** *P* < 0.0001, Student’s *t*-test (g,h); MAST Bonferonni correction (d).

To validate these findings at the protein level, we conducted an independent microbiome depletion experiment in which old mice received BrdU (100 mg/kg) at the onset of treatment to label dividing cells and track their fate after 30 days (Fig. 3E). Co-staining for BrdU and Olig2, a pan-oligodendrocyte marker, allowed identification of newly generated oligodendrocytes, while myelin basic protein (MBP) was used to assess mature myelinating cells (Fig. 3F). Quantification of BrdU+Olig2+ cells within the corpus callosum, a region with major myelinated tracts, demonstrated a significant change in newly generated oligodendrocytes following microbiome depletion (Fig. 3G), accompanied by elevated MBP expression, indicative of enhanced myelination (Fig. 3H). These results suggest that the gut microbiome exerts a regulatory influence on oligodendrocyte differentiation and myelin maintenance in the aging brain.

### Microbiome depletion enhances adult neurogenesis in aged mice

We next investigated whether reducing gut dysbiosis could influence adult hippocampal neurogenesis, a process that persists throughout life in the DG of the hippocampus and the subventricular zone (SVZ) of the lateral ventricle in mice ^26^. This form of neurogenesis is essential for learning and memory and declines sharply with age, contributing to cognitive impairment ^31^. Single-nucleus transcriptomic profiling revealed that microbiome depletion altered the transcriptional landscape of DG cells (Fig. 4A). Functional enrichment analysis of DEGs identified significant terms related to neuronal projection and dendritic development, synaptic plasticity, and higher-order functions including learning and cognition (Fig. 4B).

**Fig 4.**
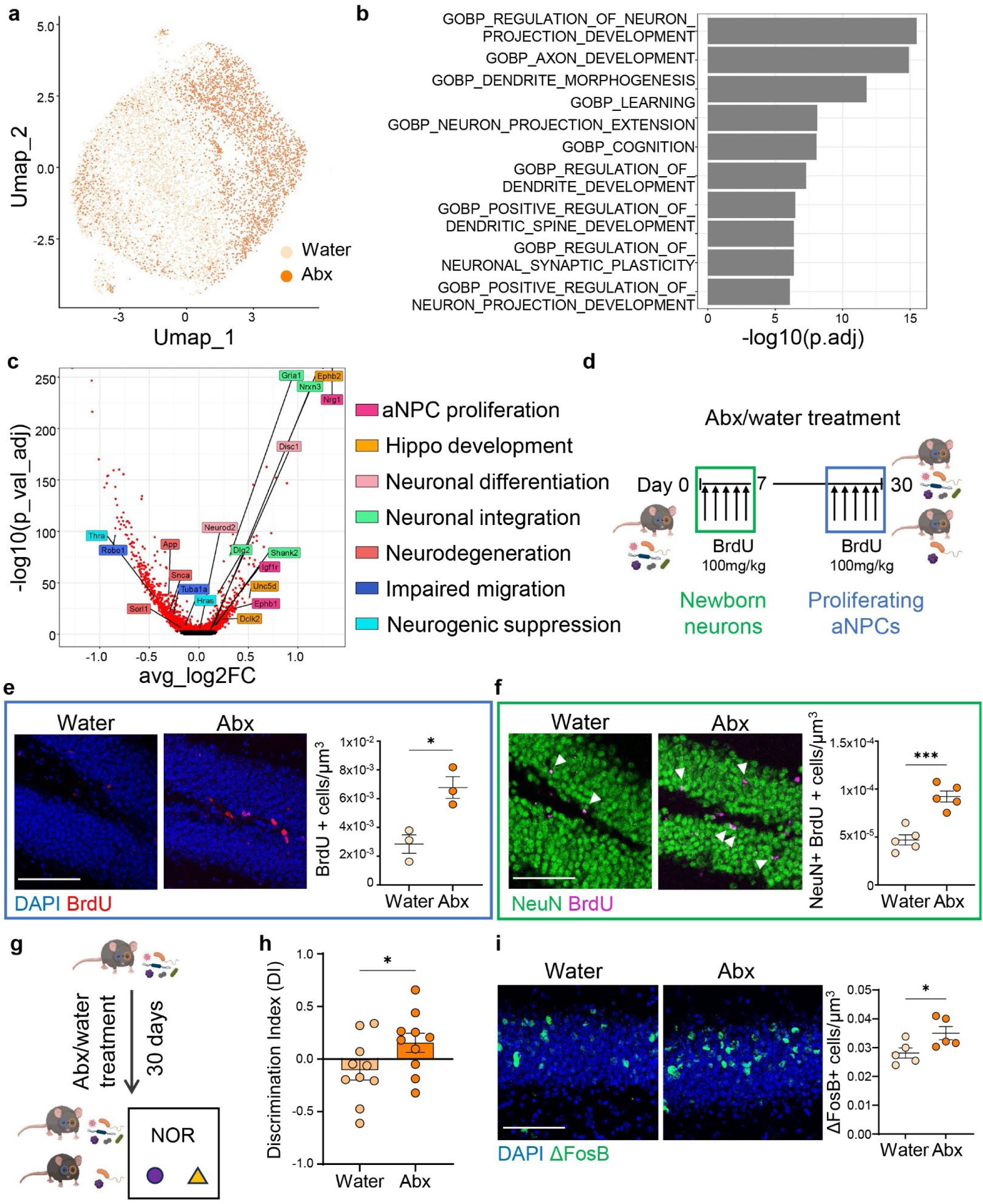
Microbiome depletion enhances adult neurogenesis in aged mice. **a,** UMAP plot showing single-nucleus transcriptomic profiles of dentate gyrus (DG) cells from aged mice, colored by treatment group. **b,** Over representation analysis of differentially expressed genes highlights pathways related to neuronal development, learning and cognition. **c,** Volcano plot of differentially expressed genes in DG cells between Abx- and water-treated mice. X-axis is the log2 foldchange, Y-axis is the log10 of the Bonferonni adjusted p-value. **d,** Schematic of BrdU administration strategies used to label newborn neurons (early BrdU, green box) and proliferating aNPCs (late BrdU, blue box) during the 30-day Abx treatment. **e,** Representative images and quantification of BrdU+ (red) cells in the DG, indicating increased aNPC proliferation in Abx-treated mice (n = 3/group). **f,** Representative images and quantification of NeuN+ (green) BrdU+ (magenta) cells in the DG, demonstrating increased generation of newborn neurons following microbiome depletion. White arrows indicate double positive cells (n = 5/group). **g,** Schematic of behavioral test design: water-treated, and Abx-treated aged mice were tested for memory using the novel object recognition test (NOR). **h,** Discrimination index in the NOR shows improved memory performance in Abx-treated mice compared to controls (n = 10/group). **i,** Representative images and quantification of ΔFosB + cells (green) in the dentate gyrus of aged mice immediately after behavioral tests (n = 5/group). Scale bar, 50 μm. All data represented as mean ± s.e.m.; \**P* < 0.05, **** *P* < 0.0001, Student’s *t*-test (d,e,g,i,l); MAST Bonferonni p-value correction (c).

Among the upregulated DEG, we observed increased expression of genes involved in neural progenitor maintenance and proliferation (Nrg1, Ephb1, Igf1r), hippocampal development and organization (Dclk2, Ephb2, Unc5d), fate commitment and neuronal differentiation (Neurod2, Disc1), and integration of adult-born neurons into existing circuits (Dlg2, Nrxn3, Shank2, Gria1). Conversely, microbiome depletion was associated with reduced expression of genes linked to neurodegeneration-associated suppression of adult neurogenesis (Snca, Sorl1, App), impaired migration and integration of newborn neurons (Robo1, Tuba1a), and suppression of progenitor proliferation and neurogenic signaling (Hrsa, Thra) (Fig. 4C, and Extended Data Fig. 3A).

To confirm changes in adult neurogenesis *in vivo*, we performed two microbiome depletion experiments in which BrdU (100 mg/kg) was administered either at the beginning or at the end of antibiotic treatment, to label newborn neurons or proliferating aNPCs, respectively (Fig. 4D). Microbiome depletion significantly increased aNPC proliferation in the aged hippocampus, as shown by the rise in BrdU+ cells within the DG (Fig. 4E). This enhanced progenitor proliferation led to a higher number of newly generated neurons, identified as NeuN+BrdU+ cells in the granule cell layer (Fig. 4F).

Given that the survival and expansion of adult neural progenitors are strongly influenced by the cellular state of the neurogenic niche, we next examined whether microbiome depletion alters senescence-associated features within the dentate gyrus. Cellular senescence has emerged as an important regulator of the adult hippocampal neurogenic niche, with senescent cells accumulating in the dentate gyrus during aging and exerting inhibitory effects on neural stem and progenitor cell function ^32^. To assess whether microbiome depletion modulates senescence-associated features of the neurogenic niche, we performed histochemical staining for senescence-associated β-galactosidase (SA-β-Gal). Notably, Abx treated mice exhibited a significant reduction in SA-β-Gal–positive cells within the subgranular zone of the dentate gyrus compared to controls (Extended Data Fig. 3B), indicating decreased senescence-associated staining in this neurogenic compartment.

Finally, to determine whether attenuation of gut dysbiosis could affect also cognitive function, we performed behavioral testing (Fig. 4G). Open field analysis revealed no significant differences between water- and Abx-treated groups in locomotion or anxiety-like behavior (Extended Data Fig. 3C-E), confirming that the treatment did not alter general activity. Strikingly, however, microbiome-depleted mice demonstrated a significantly higher discrimination index in the Novel Object Recognition (NOR) test, indicative of enhanced recognition memory (Fig. 4H). Importantly, total object exploration time was comparable across groups (Extended Data Fig. 3F), suggesting that the observed memory improvement was not due to altered motivation or attention, but rather reflected enhancement of cognitive function.

To assess neuronal activation in response to the behavioral task, we examined expression of ΔFosB in the hippocampus immediately following the behavioral tests. ΔFosB is a stable transcription factor and an indirect marker of sustained neuronal activity and plasticity ^33^. We observed increased ΔFosB immunoreactivity in the dentate gyrus of Abx-treated mice, suggesting that microbiome depletion enhances hippocampal circuit engagement during memory encoding (Fig. 4I). In addition, we detected a significant increase in the number of c-fos positive cells in the cortex of Abx-treated mice immediately following the behavioral assays (Extended Data Fig. 3G). As c-fos is an immediate early gene that reflects acute neuronal activation, this finding may indicate enhanced recruitment of cortical circuits during task performance, consistent with broader network engagement beyond the hippocampus. Collectively, these results suggest that microbiome depletion restores adult hippocampal neurogenesis, reduces cellular senescence, and enhances cognitive function in aged mice.

### Microbiome depletion attenuates microglia activation

Neuroinflammation and microglial activation are prominent hallmarks of brain aging. To assess whether microbiome depletion modulates these processes, we first examined microglial abundance and activation state. Microglia are the brain’s resident immune cells, responsible for clearing debris, pathogens, and apoptotic cells in response to injury or inflammation ^34^. In the cortex of Abx-treated aged mice, we observed a significant reduction in Iba1^+^ area and CD68 abundance compared to controls (Fig. 5A). CD68, a lysosomal glycoprotein upregulated during microglial activation, serves as a marker of phagocytic activity ^35^.These findings suggest that microbiome depletion dampens microglial activation and may attenuate neuroinflammation in the aged brain.

**Fig 5.**
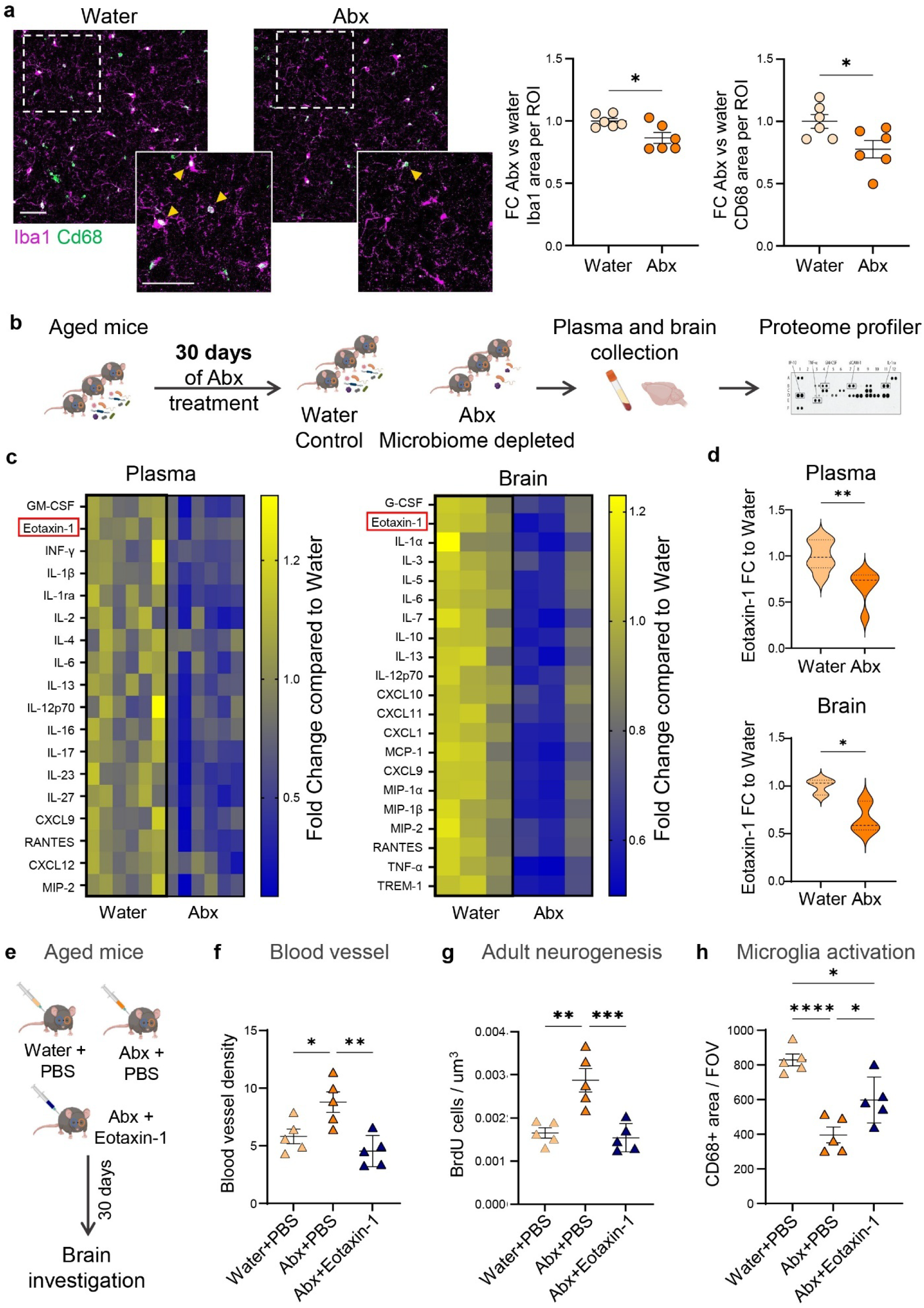
Microbiome depletion reduces age-related inflammation. **a,** Representative immunofluorescence images of Iba1+ (magenta) and CD68+ (green) microglia in the cortex of aged mice treated with water or antibiotics (Abx); quantification shows a significant reduction in both Iba1 and CD68 immunoreactivity following microbiome depletion. Yellow arrows indicate double positive cells (n = 5/group). Scale bar, 50 μm. **b,** Schematic of the experimental design: aged mice were treated with Abx or water for 30 days, after which plasma and cortical lysates were collected for cytokine profiling using a proteome array. **c,** Heatmaps showing fold change in cytokine and chemokine levels in plasma (left) and cortical lysates (right) relative to water controls. Multiple inflammatory mediators were downregulated upon Abx treatment; Eotaxin-1 (CCL11), highlighted in red, was significantly reduced in both compartments. **d,** Violin plots showing fold change in Eotaxin-1 abundance in plasma (top) and brain (bottom), confirming a significant reduction in circulating and central levels following microbiome depletion (plasma, n = 6/group; brain, n = 3/group). **e,** Schematic of rescue experiment: aged mice received antibiotics (Abx) to deplete the microbiome, in combination with either vehicle or recombinant Eotaxin-1 (n = 5/group). **f-h,** Quantification of blood vessel density (g), BrdU+ cells in the dentate gyrus (h), and CD68+ area in the cortex (i) (n = 5/group). All data represented as mean ± s.e.m.; \**P* < 0.05, ** *P* < 0.01, **** *P* < 0.0001, Student’s *t*-test (a,d); ANOVA with Tukey’s multiple-comparisons post hoc test (f-h).

### Microbiome depletion reduces age-related inflammation in multiple tissues

Given these brain-specific effects, we next asked whether microbiome depletion also exerted benefits in peripheral tissues. We focused on the retina and heart, two highly vascularized organs susceptible to age-related degeneration. In the retina, Abx-treated mice exhibited increased vascular density in the superficial layer and reduced microglial activation across all layers, as assessed by Iba1 immunoreactivity (Extended Data Fig. 4C). These effects mirror those seen in the brain, suggesting that microbiome depletion broadly reduces tissue inflammation and supports vascular remodeling. Similarly, in the heart, Masson’s trichrome staining revealed a significant decrease in ventricular fibrosis and cardiomyocyte cross-sectional area in Abx-treated mice, indicative of attenuated age-associated cardiac hypertrophy (Extended Data Fig. 4D). Together, these findings point to systemic rejuvenation following microbiome depletion and raise the possibility that factors regulated by the gut microbiome may mediate these coordinated effects across multiple organs.

To investigate how the gut might signal to the brain, especially given the effects of Abx on multiple tissues and prior studies characterizing changes in circulating factors after perturbations such as exercise and aging ^6,36^, we profiled 41 cytokines and chemokines in plasma and cortical lysates using a proteome array (Fig. 5B). Microbiome depletion led to a broad suppression of inflammatory mediators, with 18 cytokines significantly reduced in plasma, including multiple interleukins and chemokines linked to age-related inflammation ^37–39^ (Fig. 5C). A comparable reduction was observed in the cortex, where 21 cytokines were downregulated, indicating that Abx treatment also mitigates inflammation within the brain parenchyma. Notably, several of these factors, including eotaxin-1 (CCL11), RANTES (CCL5), and IL-6, can cross the blood–brain barrier, indicating a potential mechanism for peripheral immune signals to influence brain inflammation. Among the cytokines differentially abundant in both plasma and brain, seven were consistently changed, identifying them as potential systemic mediators of gut–brain communication. Of particular interest was eotaxin-1, a chemokine previously implicated in cognitive aging and neurodegeneration ^9^. Its levels were significantly decreased in both compartments following microbiome depletion (Fig. 5D). This convergence (robust downregulation, known CNS accessibility, and prior links to age-related cognitive decline) pointed to eotaxin-1 as a strong mechanistic candidate through which gut dysbiosis may shape inflammatory signaling in the aging brain.

### Eotaxin-1 mediates the effects of microbiome depletion on the aged brain

To directly test the link between systemic eotaxin-1 and brain health, we asked whether restoring circulating eotaxin-1 during microbiome depletion would be sufficient to negate the rejuvenating effects of Abx treatment. We therefore reintroduced recombinant eotaxin-1 during the Abx regimen (Fig. 5E). Restoration of circulating eotaxin-1 abolished the Abx-induced increase in blood vessel density and adult neurogenesis (Fig. 5F, G). The reduction in microglial activation observed with Abx treatment was only partially reversed by eotaxin-1 reintroduction, suggesting that additional factors or temporal dynamics may also contribute to this process (Fig. 5H).

Next, to test whether inhibition of Eotaxin-1 was sufficient to recapitulate the brain rejuvenation effects of microbiome depletion, we inhibited systemic eotaxin-1 in aged mice using a blocking antibody, as described previously ^9^ (Fig. 6A). Strikingly, eotaxin-1 inhibition phenocopied several effects of Abx treatment: treated mice exhibited improved blood vessel density (Fig. 6B), enhanced adult neurogenesis (Fig. 6C), and reduced microglial activation (Fig. 6D). Mice treated with the eotaxin-1 antibody also exhibited an increased number of oligodendrocyte precursor cells, as revealed by increased number of Pdgfra^+^ cells (Fig. 6E). In contrast to microbiome depletion, which promotes oligodendrocyte maturation and myelin production, eotaxin-1 inhibition appears to primarily expand the progenitor pool, suggesting that these interventions engage overlapping but temporally or mechanistically distinct pathways, with eotaxin-1 inhibition potentially requiring longer exposure to drive downstream myelination.

**Fig 6.**
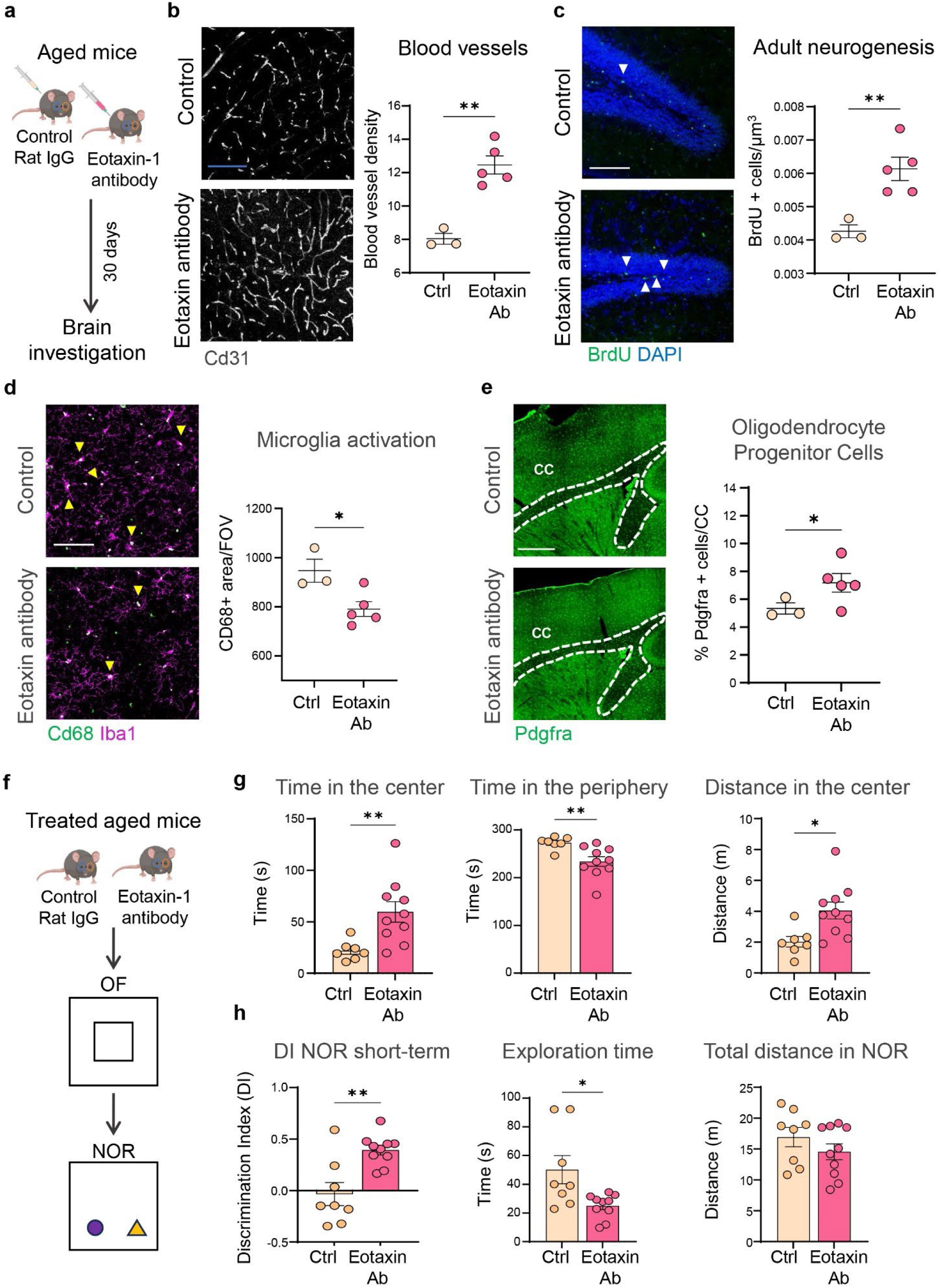
Eotaxin-1 mediates the effects of microbiome depletion on the aged brain. **a,** Schematic of the experimental design: aged mice were treated with either control IgG or an anti–eotaxin-1 neutralizing antibody for 30 days (Control, n = 3, Eotaxin Ab, n = 5). **b,** Representative CD31 immunostaining and quantification showing increased blood vessel density in the cortex of Eotaxin-1–inhibited mice. **c,** BrdU labeling (green, arrows) of proliferating cells in the DG and quantification of BrdU+ cells reveals enhanced adult neurogenesis following Eotaxin-1 inhibition. **d,** Immunofluorescence for Iba1 (magenta) and CD68 (green) shows reduced microglial activation in Eotaxin-1–treated mice. Yellow arrows indicate double positive cells. **e,** Quantification of myelinating oligodendrocyte precursor cells (Pdgfra+, green) in the corpus callosum (highlighted in white) indicates increased myelination upon Eotaxin-1 blockade. Scale bars: 100 μm (b), 50 μm (c,d), 500 μm (e). **f,** Schematic of behavioral test design: aged mice treated with Rat IgG or Eotaxin-1 antibody were tested for locomotor function and anxiety using the Open Field (OF) test, and their memory was assessed using the novel object recognition test (NOR) test. g, Time (sec) and distance spent (m) in the center or periphery of the arena during the OF test. **h,** Discrimination index, exploration time (sec) and total distance travelled (m) in the NOR test (n = 7/control; 10/Eotaxin-1 antibody). All data represented as mean ± s.e.m.; \**P* < 0.05, ** *P* < 0.01, Student’s *t*-test.

To determine whether these cellular and structural changes translate into functional improvements, we performed behavioral testing after 30 days of eotaxin-1 antibody treatment, including the open-field test and the novel-object recognition task (Fig. 6F). Remarkably, eotaxin-1 inhibition reduced anxiety-like behavior in the open-field test: treated mice spent significantly more time in the center of the arena relative to controls (Fig. 6G). Moreover, eotaxin-1 antibody treatment improved short-term memory performance in the novel-object recognition task (Fig. 6H, left), accompanied by a significant reduction in total exploration time compared to controls. Because locomotor activity was unchanged (Fig. 6H, right), reduced exploration likely reflects decreased anxiety or more efficient object investigation rather than hypoactivity.

To explore the potential source of circulating eotaxin-1, we examined gut-resident eosinophils, which are known producers of this chemokine ^40^. Flow cytometry of mouse blood showed no difference in eosinophil number upon Abx administration (Extended Data Fig. 5A). However, histological analysis revealed that eosinophil abundance in the small intestine increased with age and microbiome depletion significantly reduced eosinophil numbers (Extended Data Fig. 5B-C). Eotaxin-1 overexpression during Abx treatment led to an increase in their abundance compared to control aged mice (Extended Data Fig. 5D), suggesting that circulating Eotaxin-1 can regulate gut eosinophil levels.

Together, these findings establish eotaxin-1 as a key mediator of gut-to-brain signaling in aging. Reduction of circulating eotaxin-1 is sufficient to mimic or block the effects of microbiome depletion, supporting a microbiome–immune–brain axis that governs neurovascular and regenerative processes in the aged brain.

## Discussion

Aging is accompanied by widespread molecular, cellular, and structural changes in the brain, many of which contribute to cognitive decline and increased susceptibility to neurodegenerative diseases. Among the diverse hallmarks of brain aging are reduced vascular density, myelin degeneration, impaired neurogenesis, and chronic neuroinflammation ^3^. At the same time, emerging evidence indicates that aging is associated with a state of microbial dysbiosis, which in turn contributes to systemic inflammation and impaired brain function ^16^. In this study, we demonstrate that depletion of the gut microbiome in aged mice triggers a broad transcriptional and functional rejuvenation across multiple brain cell types. Notably, we show that antibiotic treatment leads to increased blood vessel density, stimulates the production of new myelin, and enhances the proliferation and differentiation of hippocampal neural stem cells, features commonly associated with a younger, more plastic, brain. Moreover, we observe a significant reduction in microglial reactivity, suggesting a marked attenuation of age-associated neuroinflammation. While previous studies have implicated gut microbes in neurodegeneration and cognitive decline ^41–43^, our findings position the gut microbiome as a central regulator of brain aging. By demonstrating that microbiome depletion is sufficient to reprogram multiple cellular hallmarks of aging in the brain, we highlight the gut–brain axis as a modifiable system with rejuvenation potential.

Among the emerging interventions targeting this axis, fecal microbiota transplantation (FMT) has shown promise in extending health span and lifespan in aged and progeroid mice ^44^. However, while FMT demonstrates that the microbiome can broadly influence aging trajectories, it remains a complex and poorly defined intervention with challenges in donor variability, microbial composition standardization, and clinical scalability. A key obstacle is disentangling how specific microbial taxa or their metabolites contribute to distinct aspects of brain remodeling. This task is further complicated by the high inter-individual variability in microbiome composition, even among healthy individuals, shaped by diet, environment, and host genetics ^45^. Such heterogeneity has hindered the identification of a consistent aging-associated microbial signature and complicates the development of broadly effective microbiome-based therapies.

To address this, we focused on identifying circulating mediators of the gut–brain axis that may underlie the systemic effects of microbial depletion. Communication between the gut and brain can occur via several pathways, including the vagus nerve, which senses microbial and metabolic signals and relays them to central circuits and the circulatory system, through which microbial-derived molecules, such as immune mediators and cytokines, enter the bloodstream and influence brain physiology ^46^. We found that microbiome depletion reverses age-related phenotypes not only in the brain but also in peripheral organs such as the heart and retina, supporting the hypothesis that blood-borne factors orchestrate multi-organ rejuvenation. Blood is already known to carry a number of CNS active factors whose levels are regulated by age, exercise and disease ^6^. Thus, it appears that the circulatory system also conveys at least some of the signaling molecules that the gut uses to communicate with the brain, much as it does for other aging modulators ^14^.

To identify those signaling moleules, we utilized a cytokine array. This allowed us to identify multiple known inflammatory mediators altered in both plasma and brain following Abx treatment. Among these, eotaxin-1 emerged as a compelling candidate due to its previously established association with aging and neurodegeneration ^9^. Consistent with a causal role, restoring eotaxin-1 levels during microbiome depletion was sufficient to block the rejuvenating effects of Abx treatment, preventing the increase in vascular density, neurogenesis, and microglial remodeling. Conversely, inhibition of eotaxin-1 in aged mice mimicked many of the cellular and structural benefits of microbiome depletion and additionally improved behavioral outcomes, reducing anxiety-like behavior and enhancing short-term memory. Together, these findings suggest that modulation of peripheral eotaxin-1 is a key mechanism through which the gut microbiome shapes brain aging.

We further uncovered a potential mechanistic link between eotaxin-1 and the gut immune environment. Specifically, we observed that aging is associated with an increased number of resident eosinophils in the small intestine, which parallels the age-dependent rise in circulating eotaxin-1 levels. Importantly, both eosinophil abundance and eotaxin-1 levels were normalized following microbiome depletion. Eosinophils are long-lived innate immune cells that reside in the intestinal lamina propria, where they interact closely with the epithelium and microbiota to regulate mucosal immunity. They respond to microbial signals and, in turn, secrete a range of effector molecules, including cytokines and chemokines such as eotaxin-1. Our findings suggest that age-related microbial imbalance may stimulate eotaxin-1 expression through eosinophil-driven local immune responses, thereby establishing a previously unrecognized gut–immune–brain axis with potential relevance to systemic aging and neurodegeneration.

While our study identified several cytokines altered in both plasma and brain following microbiome depletion, others, such as interferon-γ (IFN-γ), interleukin-1β (IL-1β), interleukin-23 and 17 (IL-23, IL-17), were significantly changed only in the periphery. This raises the possibility that some systemic immune signals may exert neuromodulatory effects indirectly, even in the absence of detectable changes within the brain tissue itself. Indeed, peripheral IFN-γ has been shown to influence social behavior and neural circuit function through meningeal lymphocyte activity ^47^, and IL-1β signaling has long been implicated in sickness behavior and mood regulation ^48^. Similarly, IL-23 and IL-17 have been linked to AD pathology, ALS and age-related cognitive decline ^28,37,38^. These examples suggest that blood-borne cytokines may influence brain physiology via peripheral–central signaling axes, including endothelial activation, blood–brain barrier modulation, or vagus-mediated immune sensing. Future work leveraging cell-type-specific cytokine receptor mapping and functional perturbation will be essential to dissect the contribution of peripheral cytokines to brain aging. In addition, more complete proteomic studies may reveal the existence of additional factors that direct the communication between the gut and the brain.

Together, our findings underscore the power of microbiome-derived systemic signals in coordinating age-related changes across both central and peripheral tissues. By identifying blood-borne mediators such as eotaxin-1 that act downstream of microbial composition, we highlight specific targets that could be modulated without the need for broad-spectrum antibiotics. These systemic factors may act through distinct modes; some influencing common cell types such as vascular or immune cells across tissues, while others exert more tissue-specific effects depending on receptor expression or local context. This positions the gut microbiome as a master regulator of circulating signals that orchestrate organ-specific and multi-organ aspects of aging. Dissecting the mechanisms by which these factors operate will be critical for the development of precision therapies.

This framework opens new therapeutic avenues to mitigate brain aging and potentially intervene in neurodegenerative diseases such as Alzheimer’s, where gut dysbiosis has been increasingly implicated yet mechanistic clarity remains limited ^49^. Ultimately, targeting the gut–brain axis may complement existing strategies to preserve cognitive function, offering a new frontier for promoting healthy brain aging.

## Methods

### Animals

Adult male C57BL/6J mice (≥22 months; 88–96 weeks; equivalent to 70–75 human years (Dutta & Sengupta, 2016) were obtained from the National Institute on Aging (NIA) and housed in the Harvard Biolabs Animal Facility under a 12-h light–dark cycle with ad libitum access to water and Prolab Isopro RMH 3000 chow (LabDiet). Cages contained nestlet bedding and red huts for enrichment. All procedures were approved by the Harvard University Institutional Animal Care and Use Committee (protocol AEP no. 10-23) and complied with federal and state regulations.

For antibiotic experiments, mice were cohoused in autoclaved cages with autoclaved bottled water for at least two weeks prior to dosing.

### Antibiotic administration

Microbiome depletion was performed as described ^17^ with modifications. Mice received either vehicle (water) or a four-antibiotic cocktail consisting of ampicillin sodium salt (200 mg/kg/d), neomycin trisulfate salt hydrate (200 mg/kg/d), metronidazole (200 mg/kg/d), and vancomycin hydrochloride (100 mg/kg/d; all Sigma-Aldrich) by oral gavage twice daily for 1 week. For the subsequent 2 weeks, the cocktail (vancomycin 0.5 g/l, ampicillin 1 g/l, neomycin 1 g/l) was provided in drinking water; metronidazole was excluded from the drinking water to prevent blood– brain barrier penetration ^22^. In the final week, antibiotics were administered in drinking water at the same concentrations plus once-daily oral gavage. Antibiotic-containing water was replaced every 48 h. Doses were selected based on prior reports of effective microbiome depletion ^50^. After 30 days, mice were anaesthetized with isoflurane and transcardially perfused with ice-cold PBS. Brains and peripheral organs, including small intestine, were collected for analysis. For molecular assays, hippocampus and cortex from one hemisphere were dissected and flash-frozen; the contralateral hemisphere was postfixed in 4% paraformaldehyde (PFA) overnight for histology. Fixed brains were cryoprotected in 30% sucrose (24 h), frozen, and sectioned coronally at 40 µm.

### Bacterial DNA Isolation and 16S rRNA-sequencing

Gut microbiome depletion was confirmed by 16S rRNA gene sequencing of bacterial DNA extracted from fecal samples, as previously described ^51^. Fresh fecal pellets (4–8 per mouse) were collected under sterile conditions, snap-frozen in liquid nitrogen, and stored at –80 °C until processing. Microbial DNA was isolated using the NucleoMag Pathogen kit (Macherey-Nagel, 744210.4) following the manufacturer’s protocol. DNA concentration and purity were assessed using a NanoDrop spectrophotometer, and samples (4 µg/µl) were submitted to Azenta Life Sciences for 16S rRNA amplicon sequencing. Bioinformatic analyses were performed by Azenta to profile microbial community composition, including calculation of α-diversity (within-sample diversity).

### Single-nucleus RNA-sequencing and data processing

For the snRNA-seq experiments, 4 old and 4 old antibiotic-treated (Abx) mice were analyzed, with tissue coming from both hippocampi and the isocortex. Briefly, after dissociation, cells were diluted in ice-cold PBS containing 0.4% BSA at a density of 1,000 cells/µl. For every sample, 17,400 cells were loaded into a Chromium Single Cell 3’ Chip (10x Genomics) and processed following the manufacturer’s instructions. Single-nucleus RNA-seq libraries were prepared using the Chromium v3 Single Cell 3’ Library & Gel Bead kit v2 and i7 Multiplex kit (10X Genomics). Libraries were pooled based on their molar concentrations. Pooled libraries were then loaded at 2.07 pM and sequenced on a NovaSeq instrument (Illumina) with 26 bases for read1, 91 bases for read2 and 8 bases for Index1. Cell Ranger (version 6.1.2) (10X Genomics) was used to perform sample de-multiplexing, barcode processing and single cell gene unique molecular identifier (UMI) counting, while a digital expression matrix was obtained for each experiment with default parameters, mapped to the 10x reference for mm10, version 2020-A.

#### Raw data processing and quality control for cell inclusion

Initially, there were 17,377,034 raw cells sequenced. Ambient RNA was corrected for using Cell-Bender (version 0.3.0) on the unfiltered hdf5 files, with an fpr of 0.01 over 150 epochs run on an A100 GPU, resulting in 71,939 cells post-filtering. After correction, the resulting hdf5 matrices were imported to scanpy (version 1.8.2) as adata objects and run through scrublet (v 0.2.3) to detect doublets and multiplets. The scrublet parameters were expected_doublet_rate was 0.085, sim_doublet_ratio was 2, min_counts=2, min_cells=3, min_gene_variability_pctl=75, n_prin_comps=30. Scrublet marked 323 cells as doublets or multiplets, leaving 71,616 cells.

Basic processing and visualization of the snRNA-seq data were performed using the Seurat package (version 4.3.0.900) in R (version 4.2.0). The initial dataset contained 71,616 cells with data for 29,221 genes. The average number of UMI (nCount_RNA) and non-zero genes (nFeature_RNA) are 5,562.44 and 2,182.5 respectively. Cells with greater than 12,000 UMI (nCount_RNA), less than 500 genes or greater than 5000 genes (nFeature_RNA), greater than 10% mitochondria, greater than 1.8% RPL and greater than 1% RPS were excluded from the analysis, resulting in 45,089 cells. Normalization, variable gene identification, and scaling were accomplished through the SCTransform “glmGmaPoi”, method, regressing out nFeature_RNA, percent.mito, percent.RPS, and percent.RPL. Additionally, the data were log normalized and scaled to 10,000 transcripts per cell with NormalizeData(), with variable features (n=2000) identified through FindVariableFeatures() function and scaled with ScaleData(). Next, principal component analysis (PCA) was carried out on the SCT data, and the top 40 PCs were stored. Clusters were identified with FindNeighbors() by constructing a K-nearest neighbor (KNN) graph, and clustered with the Louvain algorithm with FindClusters() at resolution 0.4 to result in 26 clusters, represented by UMAP projection.

#### Determination of cell-type-identity

We used multiple cell-specific/enriched gene markers that have been previously described in the literature to assist in determining cell-type-identity ^5,10^. Marker genes used for cluster annotation are reported in Supplementary Table 1. To further stratify the excitatory neurons, we employed SingleR (version 1.10.0) using SCT-normalized data from the Allen Brain Atlas mouse whole cortex and hippocampus SMART-seq Seurat.rda data, using the subclass labels. We identified 10 major cell types with distinct expression profiles: excitatory neurons from the hippocampus (exc_hc), neurons from the dentate gyrus (DG), inhibitory neurons (inh), excitatory neurons from the cortex (exc_ctx), astrocytes (astro), vasculature cells (vasc), oligodendrocytes (oligos), oligodendrocyte precursor cells (OPCs), immune cells (immune), and choroid plexus cells (choroid). We examined the distribution of cell type between conditions using the propeller function as adapted in Simmons 2022 ^52^. Subcluster analysis of each cell type was performed to plot transcriptional shifts in the water to antibiotic conditions.

#### Differential gene expression analysis

After initial quality control pre-processing and determination of cellular identities, we utilized the FindMarkers() function of Seurat using the MAST algorithm (version 1.22.0), with the parameters only.pos=FALSE, min.pct = 0, logfc.threshold = 0. For all mice, normalized TPM values were calculated from the RPKM and gene lengths. For single-nucleus heatmap representations, we used dittoSeq (version 1.18.0) with scaled.to.max=TRUE and Euclidean distance, ward.D2 clustering. Differentially expressed genes for each cell type are provided in Supplementary Table 2.

#### Pathway analysis

Over representation analysis (ORA) was performed via clusterProfiler (version 4.4.4) on the DEG Bonferooni adjusted p-value < 0.001 genes, with a q-value cutoff of 0.05. We used 5 gene sets: *Hallmark pathways, GO Biological Process, KEGG, BioCarta*, and *Reactome* from MSigDB (msigdbr, version 7.5.1). ORA for each cell type are summarized in Supplementary Table 3.

### Immunofluorescent Staining

Immunofluorescence Staining (IF) was conducted on 40 µm brain sections, as previously described ^53^. Sagittal brain sections (40 µm) were permeabilized in PBS containing 0.3% Triton X-100 (10 min) followed by PBS with 0.1% Triton X-100 (10 min), then blocked in 10% donkey serum in PBS with 0.1% Triton X-100 (1 h, room temperature). Sections were incubated overnight at 4 °C with primary antibodies diluted in blocking solution: goat anti-CD31 (1:100; R&D Systems, AF3628), rabbit anti-collagen IV (1:100; Bio-Rad, 2150-1470), mouse anti-ZO-1 (1:100; Thermo Fisher Scientific, 339100), mouse anti-Claudin-5 (1:100; Thermo Fisher Scientific, 352500), rat anti-CD68 (1:100; Bio-Rad, MCA1957), rabbit anti-Olig2 (Abcam, ab109186), chicken anti-MBP (Life Technologies, PA110008), rat anti-BrdU (1:100; Abcam, ab6326), mouse anti-NeuN (1:100; Millipore, MAB377), rabbit anti-Iba1 (1:100; FUJIFILM Wako, 01919741), and goat anti-Pdgfra (R&D Systems, AF1062). Nuclei were counterstained with DAPI (1:1000; Thermo Fisher Scientific, D1306). For Bromodeoxyuridine (BrdU) detection, sections were pretreated with 2 N HCl (30 min, 37 °C) before permeabilization. Following PBS washes, sections were incubated with Alexa Fluor–conjugated donkey secondary antibodies (Thermo Fisher Scientific) for 2 h at room temperature (1:500 in blocking solution), washed in PBS, air-dried, and coverslipped with ProLong mounting medium (Life Technologies, P36930). Mouse IgG (1:500; Santa Cruz Biotechnology, sc2025) served as a negative control.

Neurogenesis was quantified in the dorsal hippocampus as described ^54^. BrdU (100 mg/kg; Sigma, B5002) was administered intraperitoneally either during the first 5 days of antibiotic treatment (to label newborn neurons) or during the final 5 days before sacrifice (to label proliferating adult neural progenitor cells, aNPCs). BrdU⁺/NeuN⁺ cells were counted to determine newborn neuron numbers; BrdU⁺/DAPI⁺ cells in the subgranular zone (SGZ) were counted to quantify aNPCs.

Images were acquired on a Zeiss LSM900 confocal microscope at 10× or 20× magnification, merged into z-stacks, and processed using Fiji ^55^. Blood vessel density was quantified using AngioTool ^56^.For each mouse, 3–6 sections and at least 3 fields per region were averaged for final quantification.

Senescent cells were detected using the Senescence Cells Histochemical Staining Kit (Sigma-Aldrich, CS0030), according to the manufacturer’s instructions. Briefly, free-floating brain sections (40 µm) were mounted on slides, fixed (7 min, room temperature), rinsed in PBS, and incubated in fresh β-galactosidase staining solution (pH 4.2, 37 °C, no CO₂) for 4 h to overnight. Reactions were stopped with PBS, and coverslips were mounted with Vectashield (Vector Laboratories). Images were captured using a Nikon Eclipse 80i microscope, and the proportion of β-galactosidase–positive cells in the dentate gyrus was quantified using the Cell Counter plugin in Fiji.

### Behavioral Procedures

Mice were habituated to handling (∼30 s per day for 3 consecutive days) before testing. All assays were conducted in the light phase, with mice acclimated to the testing room for 30 min before each session. The behavioral arena was cleaned with 70% ethanol between trials.

Locomotor activity and anxiety-like behavior were assessed in a 45 × 45 cm open-field arena, as previously described ^57^. Each mouse was placed in the center of the arena and allowed to explore freely for 10 min while movement was tracked by automated software. This test also served as habituation for the novel object recognition assay. Sessions were video recorded from a camera placed below the arena, which has a glass floor (BlackBoxBio). The mouse snout, head, body, paws, tail base, and tail tip were tracked in the arena using the PalmReader DeepLabCut model (BlackBoxBio 2021 500Mar25 resnet). The average velocity of the mouse was defined as the average of the Euclidean distance the snout travelled divided by frames per second (45 fps), multiplied by 0.044 cm/pixel. The total distance the mouse travelled was defined as the cumulative Euclidean distance that the snout travelled, multiplied by 0.044 cm/pixel. The open field regions of interest (ROI) of the corners were defined as 205 pixel squares at the corners, and the center was defined as a 615 pixel square bounded by the four corners. The walls are defined as the arena space not included in the ROIs. The time spent in ROIs is measured as the time the snout spent in the ROIs, divided by frames per second (45 fps).

Recognition memory was tested as described ^58^. In the training phase, each mouse was placed in the same arena containing two identical familiar objects (FAM) for 10 min. Two hours later, one familiar object was replaced with a novel object (NEW), and mice were allowed to explore for 10 min. Object identity and location were counterbalanced across trials to avoid bias. Objects were selected based on pilot experiments confirming comparable baseline exploration times. The mouse snout, head, body, limbs, tail base, and tail tip were tracked in the arena using the PalmReader DeepLabCut model (BlackBoxBio 2021 500Mar25 resnet). Regions of interest (ROI) were defined as bounding boxes around each object. Discrimination Index (DI) was calculated the cumulative time of the snout spent in the novel object ROI, minus the cumulative time of the snout spent in the familiar object ROI, divided by frames per second (45 fps).

### Proteome Profiler to Quantify Cytokines and Chemokines in the Brain and Plasma

Relative levels of 40 cytokines and chemokines were measured in plasma and cortical lysates using the Proteome Profiler Mouse Cytokine Array Panel A (R&D Systems, ARY006), following the manufacturer’s protocol. Blood was collected by cardiac puncture into EDTA-coated tubes, centrifuged at 2,000 × g for 15 min, and 100 µl of plasma was used per array. For cortical lysates, tissue was homogenized in PBS containing protease inhibitors, supplemented with 1% Triton X-100, frozen at –80 °C overnight, thawed, and centrifuged at 10,000 × g for 5 min to remove debris. Protein concentrations were determined by Bradford assay, and 500 µg of total protein was loaded per membrane. Dot blots were developed using SuperSignal™ West Pico PLUS chemiluminescent substrate (Thermo Fisher Scientific, 34580) and imaged in a darkroom. Signal intensities were quantified using ImageJ (NIH) according to the Dot Blot Analysis Protocol, with normalization to internal reference spots.

### Eotaxin-1 manipulation

For depletion experiments, 23-month-old C57BL/6J mice received intraperitoneal injections of an anti–Eotaxin-1 antibody (50 ng/g body weight; R&D Systems, 42285) every 3 days for 30 days. An isotype-matched rat IgG2a (50 ng/g body weight; R&D Systems, 54447) was used as control.

For overexpression experiments, a separate cohort of 23-month-old C57BL/6J mice was treated with antibiotics for 30 days (as above) and concurrently injected intraperitoneally with recombinant mouse Eotaxin (10 ng/g body weight in PBS (2 μg/ml stock; R&D Systems, 420ME020) every 3 days. Control groups included antibiotic-treated mice injected with PBS and water-treated mice injected with PBS (5 µl/g body weight).

At the end of each protocol, mice were perfused, and brains and peripheral organs were collected and processed as described above.

### Retina whole mount and staining

Retinal flat mounts were prepared as described ^59^. Eyes were fixed in 4% PFA for 1 h at room temperature, the anterior capsule was removed, and the lens nuclei and cortex were dissected away. The retina was separated from the choroid and sclera, cleared of adherent vitreous with fine brushes, and cut into four leaflets. Samples were blocked in 10% donkey serum with 0.1% Triton X-100 in PBS for 2 h at 4 °C, then incubated overnight at 4 °C with primary antibodies diluted in blocking buffer. Endothelial cells were labeled with fluorescent isolectin GS-IB4 (Thermo Fisher Scientific, I21412), and microglia were labeled with Iba1 antibody as in brain staining protocols. After PBS washes, samples were incubated for 2 h at room temperature with Alexa Fluor conjugated secondary antibodies (Thermo Fisher Scientific), counterstained with DAPI, and mounted in ProLong Antifade medium. Images were acquired on a Zeiss LSM900 confocal microscope (10× or 20×). Blood vessel density was quantified in deep, intermediate, and superficial vascular plexuses using AngioTool, and microglial abundance was measured in z-stacks using Fiji.

### Analysis of blood cell composition

Blood was collected by cardiac puncture as described above. Complete blood counts, including white blood cell, red blood cell, hemoglobin, hematocrit, mean corpuscular volume, and platelet levels, were obtained using an Element HT5 hematology analyzer (Heska).

### Heart analysis

Hearts were collected and postfixed in 4% paraformaldehyde (PFA) overnight for histology. Fixed hearts were embedded in paraffin and sectioned transversely at 8 µm until further analysis. and sectioned transversely at the mid-ventricular level

#### Fibrosis analysis

Masson trichrome staining was performed on paraffin-embedded heart sections. Ventricular sections were deparaffinized in Histoclear (National Diagnostics) until no paraffin granules were visible, followed by graded ethanol dilutions (100%, 95%, 70%, H2O) to rehydrate tissue samples. Slides were immersed in Bouin’s solution (Sigma-Aldrich) overnight at room temperature, then incubated in Weigert’s iron hematoxylin (scarlet–acid red solution; Sigma-Aldrich) for 10 min. The sections were then immersed in phosphotungstic acid for 5 min and immediately transferred into aniline blue solution for 15 min. Slides were quickly dipped in 1% acetic acid before subsequent dehydration and coverslipped with Dako mounting medium.

In addition, fibroblast activation was assessed by immunostaining using α-smooth muscle actin (α-SMA). Sections were deparaffinized in Histoclear and rehydrated through graded ethanol dilutions (100%, 95%, 70%) as described above. After washing in PBS, sections were permeabilized with 0.5% Triton X-100 in PBS for 10 min and blocked in 10% donkey serum for 1 h at room temperature. Sections were incubated overnight at 4 °C with primary antibody against α-SMA (1:100; Cell Signaling) diluted in blocking buffer. Following PBS washes, sections were incubated with Alexa Fluor–conjugated secondary antibodies (Thermo Fisher Scientific) for 1 h at room temperature (1:1000 in blocking solution) with Hoesch (1:10 000), washed in PBS, and coverslipped with Dako mountung medium.

Images were acquired on the AxioScan Z.1 (Zeiss) at 20× magnification. Collagen deposition was quantified as the percentage of aniline blue–positive area over total tissue area using Fiji software. Fibroblast activation was expressed as the percentage of α-SMA–positive area over total myocardial area, calculated using Fiji thresholding. For each mouse, three transverse sections from comparable mid-ventricular levels were analyzed, and the mean value was used for statistical analysis.

#### Cross-sectional area

To assess cardiomyocyte cross-sectional area, the heart sections were processed as described above and stained by immunostaining using wheat germ agglutinin (WGA)-647 (ThermoFisher Scientific) overnight at 4 °C. Following PBS washes, sections were incubated with Hoesch (1:10 000), washed in PBS, and coverslipped with Dako medium.

Slides were imaged using a Zeiss Axio Scan Z.1 slide scanner with a 20× objective. For crosssectional area analysis, cardiomyocytes with circular profiles and central nuclei were measured in Fiji from WGA-stained sections, and at least 100 myocytes per heart were quantified. Fibroblast activation was expressed as the percentage of α-SMA–positive area over total myocardial area, calculated using Fiji thresholding. For each mouse, three transverse sections from equivalent ventricular regions were analyzed and averaged for statistical comparisons.

### Intestine analysis

Small intestines were isolated from mice, flushed thoroughly with PBS, post-fixed in 4% PFA overnight at 4 °C, and transferred to 70% ethanol. Tissues were rolled into “Swiss rolls,” placed into cassettes, and processed for paraffin embedding. Paraffin blocks were sectioned at 5 µm using a microtome, and sections were mounted on Superfrost slides. For staining, slides were deparaffinized in Histoclear (2 × 10 min), rehydrated through graded ethanols (100% ethanol 2 × 10 min, 95% ethanol 5 min, 70% ethanol 5 min), rinsed in deionized water, and equilibrated in PBS for 10 min followed by incubation in 0.1% PBST for 10 min. Slides were then outlined with a hydrophobic barrier and incubated in blocking buffer (1% donkey serum in 0.1% PBST) for 30 min at room temperature. Primary antibodies were diluted according to manufacturer recommendations and applied overnight at 4 °C. Conditions were optimized as needed for tissue and antibody performance. For eosinophil staining, we used a mouse anti–Siglec-F/CD170 antibody (Life Technologies # 14170282). The following day, slides were washed in 0.1% PBST and incubated with the appropriate fluorescent secondary antibody (1:1000) for 60 min at room temperature, protected from light. After additional PBST and PBS washes, sections were counterstained with DAPI (1:1000), rinsed in PBS, and mounted using ProLong antifade medium. Fluorescent images were acquired using a standard epifluorescence microscope.

### Statistics

Data are presented as mean ± SEM. Statistical analyses were performed in GraphPad Prism 10 (GraphPad Software). Comparisons between two groups were made using unpaired, two-tailed Student’s t-tests. Comparisons among three or more groups were performed using one-way ANOVA followed by Tuckey’s post hoc test. Exact sample sizes (n) are reported in figure legends and refer to biological replicates (individual mice). Exact statistical model details and P values are provided in Supplementary Table 4. Differences were considered statistically significant at P < 0.05. No statistical methods were used to predetermine sample size. Variance was assumed to be equal and not formally tested, and data was assumed to be normal but not formally tested.

## Data and materials availability

All data supporting the findings of this study are available within the paper and its Supplementary Information or from the corresponding author upon reasonable request. Data exploration of this snRNA-seq study is currently available at https://rubinlab.connect.hms.harvard.edu/microbiome/. Raw sequencing data will be deposited to GEO and the processed counts matrices to the Broad Single Cell Portal upon acceptance of this manuscript.

## Acknowledgements

We thank the members of the Rubin laboratory for helpful discussions and advice in different aspects of the study. We thank J. LaLonde and I. Adatto for their technical support. We also thank the staff members of the Harvard Biolabs Animal Facility, the Harvard Center for Biological Imaging and the Harvard Stem Cell and Regenerative Biology Histology Core for their continuous support and assistance. Experimental schematics were created using BioRender under an institutional license with full publication rights (Harvard University). This work was supported by a generous gift from the Vranos Family Foundation (L.L.R.), a grant from the National Institutes of Health (NIH)/National Institute of Aging (grant no. 1R01AG072086) (L.L.R.), and the Simons Foundation (Collaboration on Plasticity and the Aging Brain) (L.L.R.). The funders had no role in the study design, experiments performed, data collection, data analysis and interpretation, or preparation of the manuscript. However, we would like to thank Michael Vranos for his ongoing interest in our work and for stimulating discussions concerning the gut-brain axis.

## Contributions

C.G. and L.L.R. conceived and designed the study. C.G. performed the microbiome depletion, eotaxin manipulation, and behavioral experiments. M.J., C.C.D., and K.K. assisted with antibiotic administration and mouse handling. C.G. and F.L. performed the scRNA-seq experiments. K.M.H. and F.L. processed the scRNA-seq data, and K.M.H. developed the computational framework and performed all associated analyses. C.G. interpreted the data. C.G., M.J., C.C.D., K.M.W., S.P., and R.M.G. performed staining experiments and quantification. L.B.D. analyzed the heart samples. C.G. and G.W. conducted the retinal analyses. A.K. performed the blood cell analyses. C.G. designed and/or performed validation experiments. K.M.H. and Q.X. built the online portal. C.G. supervised aspects of the study. L.L.R. directed the study and secured funding. C.G. wrote the original draft of the manuscript. K.M.H., F.L., K.K., K.M.W., R.M.G., L.B.D., R.T., and L.L.R. provided critical feedback and edited the manuscript. All authors reviewed the manuscript and approved its submission.

## Corresponding authors

Correspondence to.

## Extended data

**Extended data Fig. 1.**
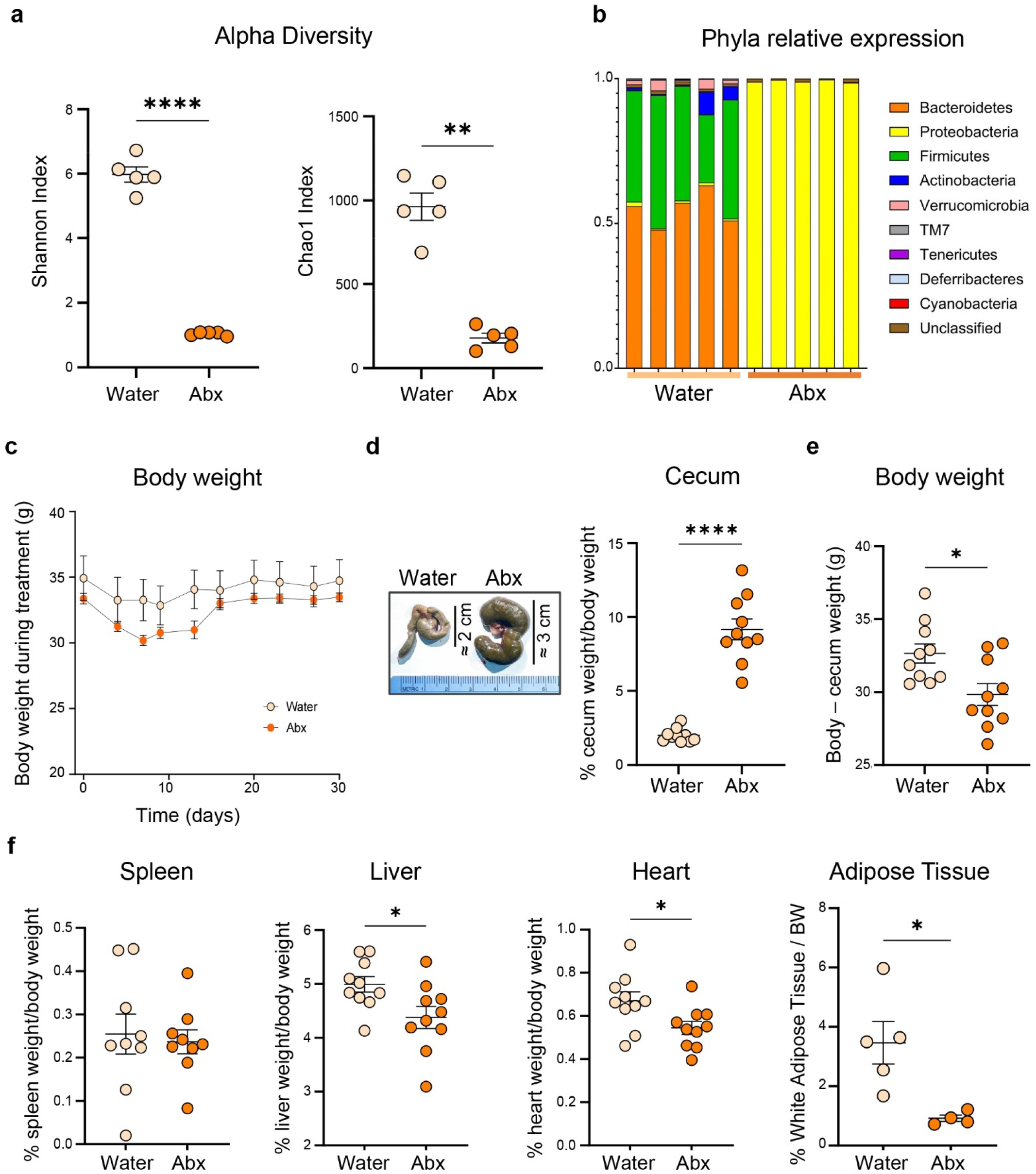
Antibiotic treatment alters organ weights and gut microbiome composition. **a,** Microbial α-diversity represented as Shannon and Chao1 index in fecal samples, assessed by 16S rRNA sequencing after Abx treatment. **b,** Relative abundance of bacterial phyla in fecal material following microbiome depletion. **c,** Body weights of aged mice monitored daily during the 30-day water or antibiotic (Abx) treatment. **d,** Representative images of ceca (left) and quantification of cecum weight as a percentage of total body weight following treatment. **e,** Body weight adjusted for cecum mass (total body weight minus cecum weight). **f,** Relative weights of selected peripheral organs (liver, heart, white adipose tissue) normalized to body weight. All data represented as mean ± s.e.m.; \**P* < 0.05, ** *P* < 0.01, **** *P* < 0.0001, Student’s *t*-test.

**Extended data Fig. 2.**
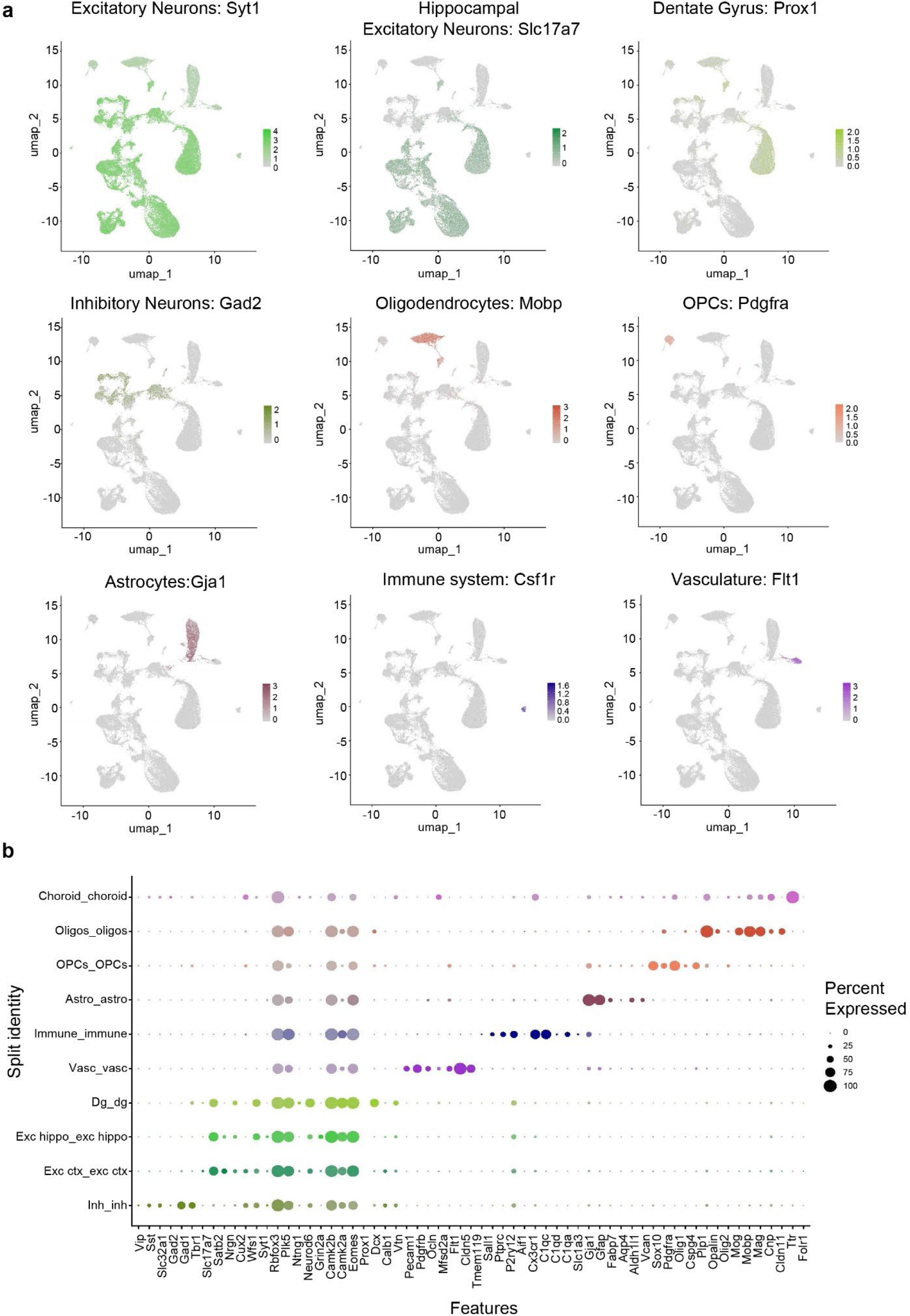
Marker gene expression used to annotate single-nucleus RNA-seq clusters. **a,** Feature plots showing the expression of representative marker genes used to identify major brain cell types in the UMAP embedding of single-nucleus transcriptomes. Genes include Slc17a7 and Syt1 (excitatory neurons), Prox1 (dentate gyrus granule neurons), Gad2 (inhibitory neurons), Mobp (mature oligodendrocytes), Pdgfra (oligodendrocyte precursor cells), Gja1 (astrocytes), Csf1r (microglia), and Flt1 (vascular endothelial cells). Expression intensity is color-coded per cell across the UMAP space. **b,** Dotplot of marker genes used to annotate major cell clusters. Dot size represents the percentage of cells expressing each marker, and dot color indicates average expression level.

**Extended data Fig. 3.**
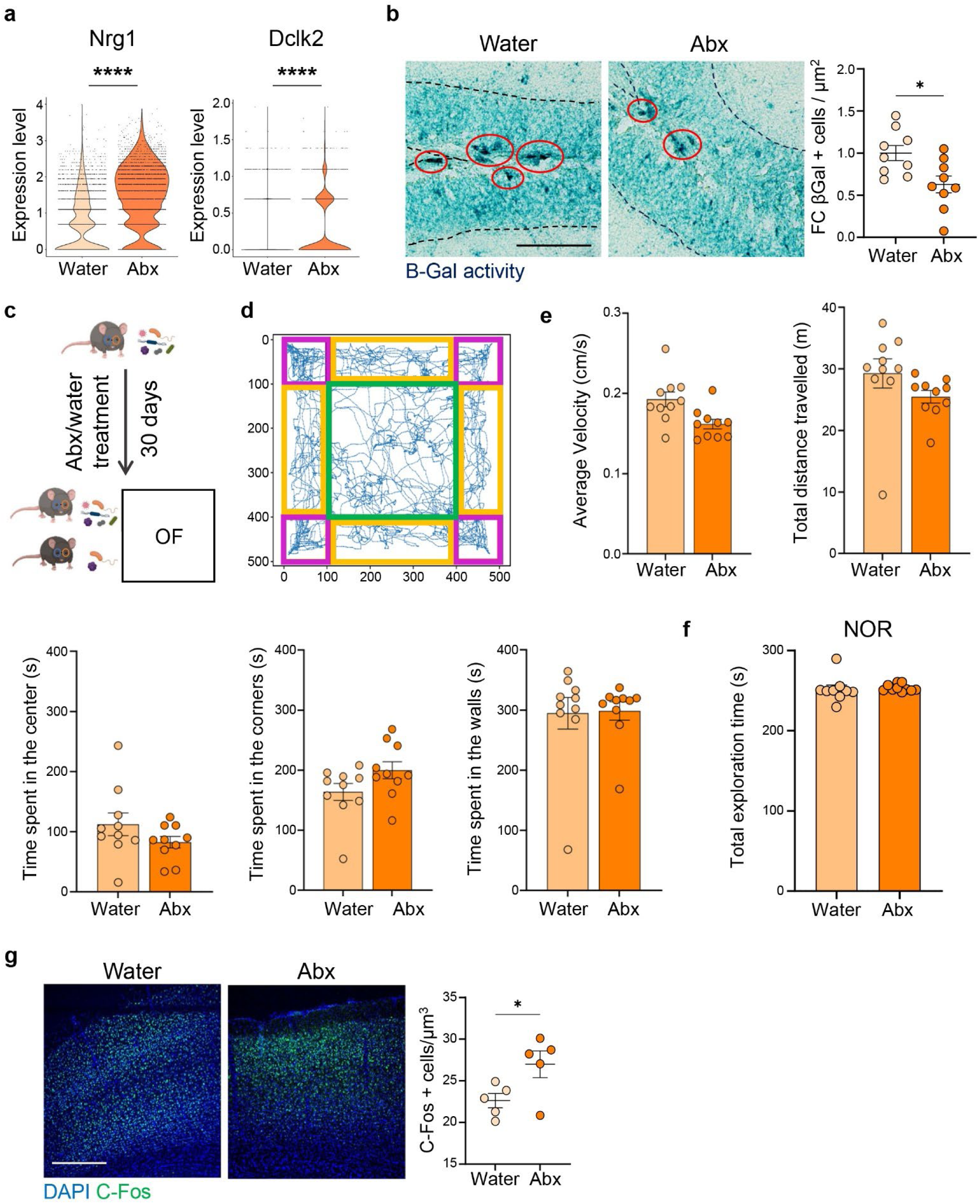
Microbiome depletion reduces hippocampal senescence, and it does not affect anxiety and locomotor activity of aged mice. **a,** Violin plots showing increased expression of Nrg1 and Dclk2, genes associated with adult neural progenitor cell (aNPC) proliferation and hippocampal organization, in Abx-treated mice. **b,** Representative images of senescence-associated β-galactosidase (SA-β-Gal) staining and quantification of β-Gal+ cells in the DG, revealing decreased cellular senescence upon microbiome depletion (n = 9/group). **c,** Scheme of behavior experiment. **d,** Representative movement trace from the open field test showing center (green), corners (magenta) and walls (yellow) zones. **e,** Quantification of average velocity, total distance travelled, and time spent in the center, corners, or walls during the open field test; no significant differences were observed between water- and Abx-treated mice (n = 10/group). **f,** Total object exploration time during the novel object recognition test was unchanged between groups, indicating similar exploratory drive and attention (n = 10/group). **g,** Representative images and quantification of c-Fos + cells (green) in the prefrontal cortex of aged mice immediately after behavioral tests (n = 5/group). Scale bar, 50 μm (b) and 500 μm (g). All data represented as mean ± s.e.m.; \**P* < 0.05, **** *P* < 0.0001, Student’s *t*-test (d,e,g,i,l); MAST Bonferonni p-value correction (b).

**Extended data fig 4.**
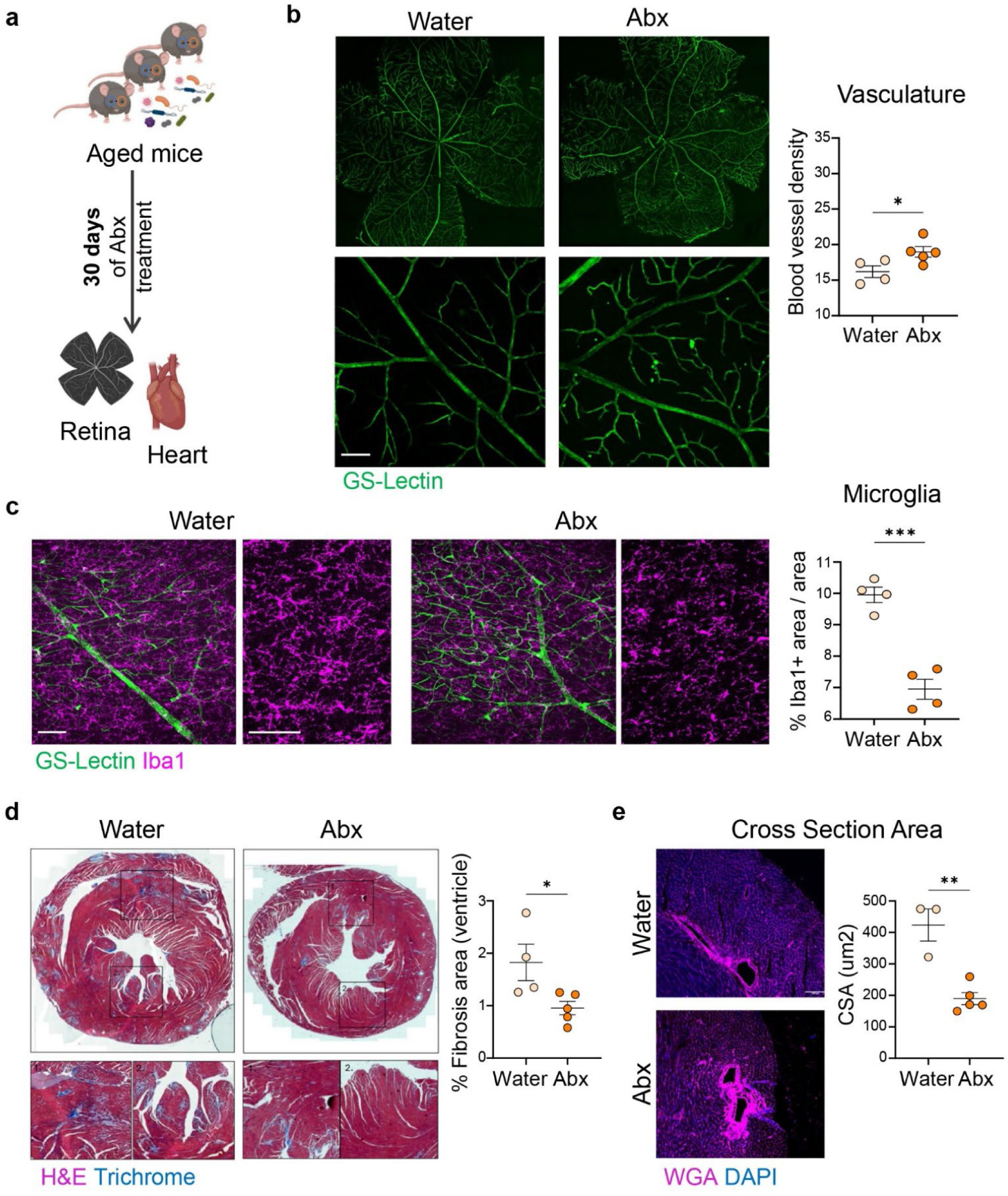
Microbiome depletion effects extend beyond the brain. **a,** Scheme of the experiment. **b,** Quantification of blood vessel density in the superficial layer of the retina via whole mount staining with GS-Lectin (green). **c,** Quantification of Iba1 area (magenta) in the three layers of the retina. **d,** Quantification of fibrosis (collagen deposition) in heart sections via Masson trichrome staining, quantified as the percentage of aniline blue–positive area over left ventricle area. **e,** Quantification of cardiomyocyte cross-sectional area, detected by wheat germ agglutinin (WGA) staining (magenta). Cardiomyocytes with circular profiles were measured; at least 100 myocytes per heart were quantified. Scale bars 100 µm. All data represented as mean ± s.e.m.; \**P* < 0.05, ** *P* < 0.01, *** *P* < 0.001, Student’s *t*-test.

**Extended data Fig 5.**
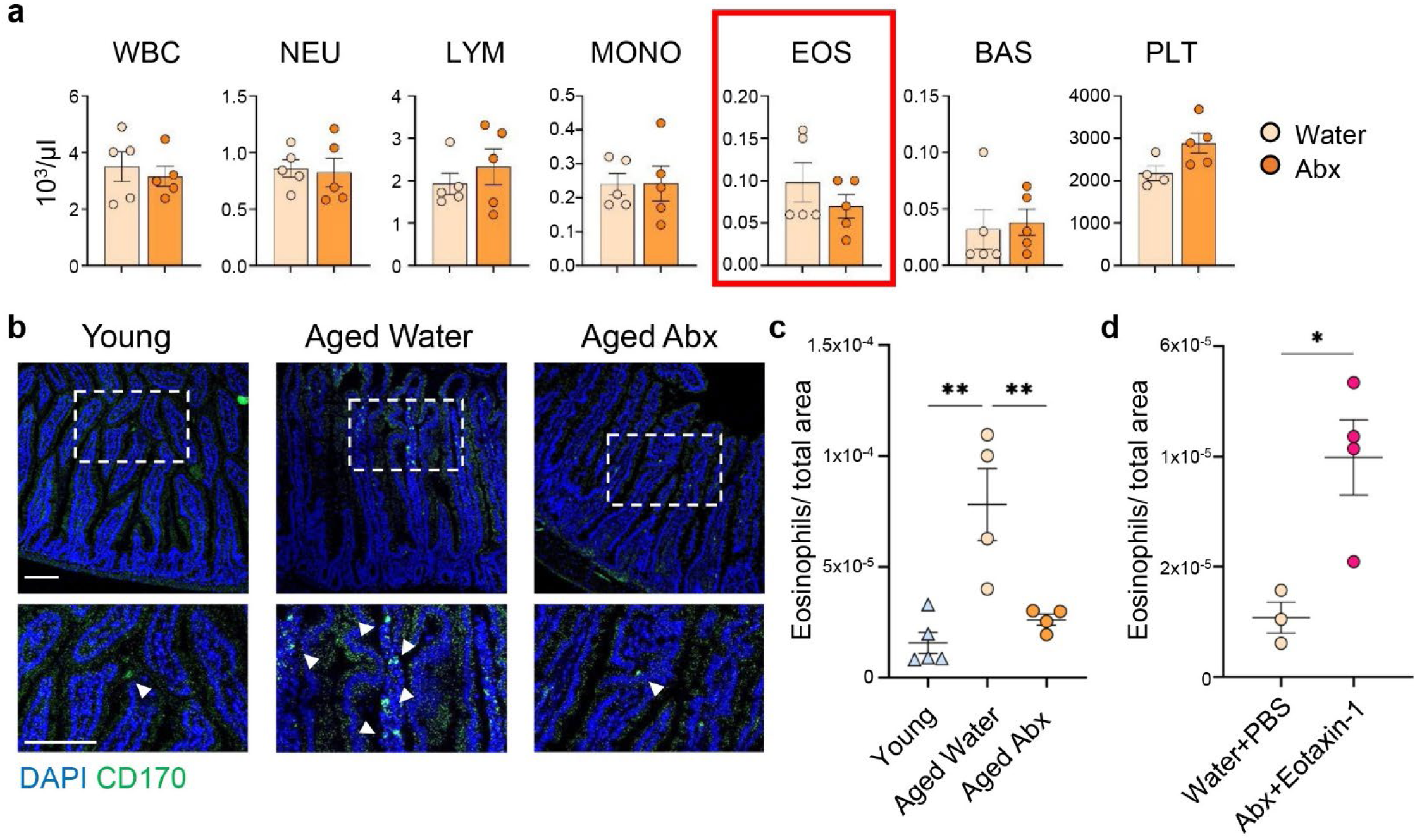
Intestinal eosinophils increase with age and may contribute to systemic Eotaxin-1 levels. **a,** Flow cytometry analysis of peripheral blood from aged mice showing the frequency of major immune cell populations, including eosinophils, following Abx treatment. **b,** Representative images of CD170+ eosinophils (green) in the intestinal epithelium of young, aged control, and aged Abx-treated mice. **c,** Quantification of CD170+ eosinophils normalized to intestinal villus area in young, aged control, and aged Abx-treated mice (n = 4/group). **d,** Quantification of CD170+ eosinophils in the intestinal villi of aged mice treated with water + PBS or Abx + Eotaxin-1 (n = 3 control, n = 4 Abx + Eotaxin). White arrows indicate CD170 positive cells. Scale bar 100 µm. All data represented as mean ± s.e.m.; \**P* < 0.05, ** *P* < 0.01. ANOVA with Tukey’s multiple-comparisons post hoc test (c), Student’s *t*-test (d).

## Supplementary information

**Supplementary Table 1. Marker genes used for cluster annotation**

List of canonical marker genes used to assign cell-type identities across all clusters in the snRNA-seq dataset.

**Supplementary Table 2. Differential gene expression for each cell type**

MAST differential gene expression analysis with unadjusted p-values (p_val), average log2 foldchange (avg_log2FC), percent expression of the gene within that cluster (pct.1), percent expression of the gene in all other clusters (pct.2), and Bonferonni-adjusted p-value (p_val_adj).

**Supplementary Table 3. Over representation analysis (ORA) across cell types**

Over representation analysis for each cell type, based on MAST adjusted p-values < 0.001 DEG profiles. The MSigDB pathway (ID, description), Number of genes in the DEG over the number of genes in the gene set (Gene Ratio), number of genes compared to the background (BgRatio), Fisher’s exact test p-value (pvalue), adjustd p-value (p.adjust), Benjamini-Hochberg FDR (qvalue), genes in the gene set (geneID), and number of DEG genes in the gene set (Count).

**Supplementary Table 4. Statistical test and *P* values**

## References

1. World Health Organization. Ageing and health. https://www.who.int/news-room/fact-sheets/detail/ageing-and-health https://www.who.int/news-room/fact-sheets/detail/ageing-and-health (2024).

2. Hou, Y. et al. Ageing as a risk factor for neurodegenerative disease. Nat. Rev. Neurol. 15, 565–581 (2019).

3. López-Otín, C., Blasco, M. A., Partridge, L., Serrano, M. & Kroemer, G. Hallmarks of aging: An expanding universe. Cell 186, 243–278 (2023).

4. Tartiere, A. G., Freije, J. M. P. & López-Otín, C. The hallmarks of aging as a conceptual framework for health and longevity research. Frontiers in Aging 5, 1334261 (2024).

5. Ximerakis, M. et al. Single-cell transcriptomic profiling of the aging mouse brain. Nature Neuroscience 2019 22:10 22, 1696–1708 (2019).

6. Bieri, G., Schroer, A. B. & Villeda, S. A. Blood-to-brain communication in aging and rejuvenation. Nat. Neurosci. 26, 379–393 (2023).

7. Gildea, H. K. & Liddelow, S. A. Mechanisms of astrocyte aging in reactivity and disease. Mol. Neurodegener. 20, 21 (2025).

8. Bouchard, J. & Villeda, S. A. Aging and brain rejuvenation as systemic events. J. Neurochem. 132, 5–19 (2015).

9. Villeda, S. A. et al. The ageing systemic milieu negatively regulates neurogenesis and cognitive function. Nature 2011 477:7362 477, 90–94 (2011).

10. Ximerakis, M. et al. Heterochronic parabiosis reprograms the mouse brain transcriptome by shifting aging signatures in multiple cell types. Nature Aging 2023 3:3 3, 327–345 (2023).

11. Horowitz, A. M. et al. Blood factors transfer beneficial effects of exercise on neurogenesis and cognition to the aged brain. Science 369, 167 (2020).

12. Katsimpardi, L. et al. Vascular and neurogenic rejuvenation of the aging mouse brain by young systemic factors. Science (1979). 344, 630–634 (2014).

13. Schroer, A. B. et al. Platelet factors attenuate inflammation and rescue cognition in ageing. Nature 620, 1071–1079 (2023).

14. De Miguel, Z. et al. Exercise plasma boosts memory and dampens brain inflammation via clusterin. Nature 600, 494–499 (2021).

15. Villeda, S. A. et al. Young blood reverses age-related impairments in cognitive function and synaptic plasticity in mice. Nat. Med. 20, 659–663 (2014).

16. Ghosh, T. S., Shanahan, F. & O’Toole, P. W. The gut microbiome as a modulator of healthy ageing. Nat. Rev. Gastroenterol. Hepatol. 19, 565–584 (2022).

17. Burberry, A. et al. C9orf72 suppresses systemic and neural inflammation induced by gut bacteria. Nature 2020 582:7810 582, 89–94 (2020).

18. Harach, T. et al. Reduction of Abeta amyloid pathology in APPPS1 transgenic mice in the absence of gut microbiota. Scientific Reports 2017 7:1 7, 41802-(2017).

19. Minter, M. R. et al. Antibiotic-induced perturbations in gut microbial diversity influences neuro-inflammation and amyloidosis in a murine model of Alzheimer’s disease. Scientific Reports 2016 6:1 6, 30028-(2016).

20. Zarrinpar, A. et al. Antibiotic-induced microbiome depletion alters metabolic homeostasis by affecting gut signaling and colonic metabolism. Nature Communications 2018 9:1 9, 1–13 (2018).

21. Tan, J. et al. Evaluation of an Antibiotic Cocktail for Fecal Microbiota Transplantation in Mouse. Front. Nutr. 9, 918098 (2022).

22. Lupori, L. et al. The gut microbiota of environmentally enriched mice regulates visual cortical plasticity Graphical abstract. https://doi.org/10.1016/j.celrep.2021.110212 (2022) doi:10.1016/j.celrep.2021.110212.

23. Liu, M. N., Lan, Q., Wu, H. & Qiu, C. W. Rejuvenation of young blood on aging organs: Effects, circulating factors, and mechanisms. Heliyon 10, e32652 (2024).

24. Brandhorst, S. et al. A Periodic Diet that Mimics Fasting Promotes Multi-System Regeneration, Enhanced Cognitive Performance, and Healthspan. Cell Metab. 22, 86–99 (2015).

25. Zarrinpar, A. et al. Antibiotic-induced microbiome depletion alters metabolic homeostasis by affecting gut signaling and colonic metabolism. Nature Communications 2018 9:1 9, 1–13 (2018).

26. Boldrini, M. et al. Human Hippocampal Neurogenesis Persists throughout Aging. Cell Stem Cell 22, 589–599.e5 (2018).

27. Fang, R. et al. Conservation and divergence of cortical cell organization in human and mouse revealed by MERFISH. Science (1979). 377, (2022).

28. Limone, F. et al. Myeloid and lymphoid expression of C9orf72 regulates IL-17A signaling in mice. Sci. Transl. Med. 16, 31 (2024).

29. Yao, Z. et al. A high-resolution transcriptomic and spatial atlas of cell types in the whole mouse brain. Nature 2023 624:7991 624, 317–332 (2023).

30. Santisteban, M. M. & Iadecola, C. The pathobiology of neurovascular aging. Neuron 113, 49–70 (2025).

31. Gonçalves, J. T., Schafer, S. T. & Gage, F. H. Adult Neurogenesis in the Hippocampus: From Stem Cells to Behavior. Cell 167, 897–914 (2016).

32. Fatt, M. P. et al. Restoration of hippocampal neural precursor function by ablation of senescent cells in the aging stem cell niche. Stem Cell Reports 17, 259 (2022).

33. Nestler, E. J. ΔFosB: a transcriptional regulator of stress and antidepressant responses. Eur. J. Pharmacol. 753, 66 (2014).

34. Colonna, M. & Butovsky, O. Microglia Function in the Central Nervous System During Health and Neurodegeneration. Annu. Rev. Immunol. 35, 441 (2017).

35. Helmut, K., Hanisch, U. K., Noda, M. & Verkhratsky, A. Physiology of microglia. Physiol. Rev. 91, 461–553 (2011).

36. De Miguel, Z. et al. Exercise plasma boosts memory and dampens brain inflammation via clusterin. Nature 2021 600:7889 600, 494–499 (2021).

37. Brigas, H. C. et al. IL-17 triggers the onset of cognitive and synaptic deficits in early stages of Alzheimer’s disease. Cell Rep. 36, (2021).

38. Mohammadi Shahrokhi, V., et al. IL-17A and IL-23: plausible risk factors to induce age-associated inflammation in Alzheimer’s disease. Immunol. Invest. 47, 812–822 (2018).

39. Nitsch, L., Schneider, L., Zimmermann, J. & Müller, M. Microglia-Derived Interleukin 23: A Crucial Cytokine in Alzheimer’s Disease? Front. Neurol. 12, 639353 (2021).

40. Teixeira, A. L., Gama, C. S., Rocha, N. P. & Teixeira, M. M. Revisiting the role of eotaxin-1/CCL11 in psychiatric disorders. Front. Psychiatry 9, 370423 (2018).

41. Sampson, T. R. et al. Gut Microbiota Regulate Motor Deficits and Neuroinflammation in a Model of Parkinson’s Disease. Cell 167, 1469–1480.e12 (2016).

42. Loh, J. S. et al. Microbiota–gut–brain axis and its therapeutic applications in neurodegenerative diseases. Signal Transduct. Target. Ther. 9, 1–53 (2024).

43. Chandra, S., Sisodia, S. S. & Vassar, R. J. The gut microbiome in Alzheimer’s disease: what we know and what remains to be explored. Mol. Neurodegener. 18, 1–21 (2023).

44. Bárcena, C. et al. Healthspan and lifespan extension by fecal microbiota transplantation into progeroid mice. Nature Medicine 2019 25:8 25, 1234–1242 (2019).

45. Huttenhower, C. et al. Structure, function and diversity of the healthy human microbiome. Nature 2012 486:7402 486, 207–214 (2012).

46. Morais, L. H., Schreiber, H. L. & Mazmanian, S. K. The gut microbiota–brain axis in behaviour and brain disorders. Nat. Rev. Microbiol. 19, 241–255 (2021).

47. Filiano, A. J. et al. Unexpected role of interferon-γ 3 in regulating neuronal connectivity and social behaviour. Nature 535, 425–429 (2016).

48. Dantzer, R., O’Connor, J. C., Freund, G. G., Johnson, R. W. & Kelley, K. W. From inflammation to sickness and depression: When the immune system subjugates the brain. Nat. Rev. Neurosci. 9, 46–56 (2008).

49. Nystuen, K. L. et al. Alzheimer’s Disease: Models and Molecular Mechanisms Informing Disease and Treatments. Bioengineering 11, 45 (2024).

50. Tirelle, P. et al. Comparison of different modes of antibiotic delivery on gut microbiota depletion efficiency and body composition in mouse. BMC Microbiol. 20, (2020).

51. Love, M. I., Huber, W. & Anders, S. Moderated estimation of fold change and dispersion for RNA-seq data with DESeq2. Genome Biol. 15, (2014).

52. Simmons, S. Cell Type Composition Analysis: Comparison of statistical methods. bioRxiv 2022.02.04.479123 (2022) doi:10.1101/2022.02.04.479123.

53. Gasperini, C. et al. Piwil2 (Mili) sustains neurogenesis and prevents cellular senescence in the postnatal hippocampus. EMBO Rep. 24, (2023).

54. Pons-Espinal, M., et al. MiR-135a-5p Is Critical for Exercise-Induced Adult Neurogenesis. Stem Cell Reports 12, (2019).

55. Schindelin, J. et al. Fiji: An open-source platform for biological-image analysis. Nat. Methods 9, 676–682 (2012).

56. Zudaire, E., Gambardella, L., Kurcz, C. & Vermeren, S. A computational tool for quantitative analysis of vascular networks. PLoS One 6, (2011).

57. Seibenhener, M. L. & Wooten, M. C. Use of the Open Field Maze to Measure Locomotor and Anxiety-like Behavior in Mice. J. Vis. Exp. 52434 (2015) doi:10.3791/52434.

58. Denninger, J. K., Smith, B. M. & Kirby, E. D. Novel Object Recognition and Object Location Behavioral Testing in Mice on a Budget. J. Vis. Exp. 2018, 10.3791/58593 (2018).

59. Marra, K. V., et al. Bioactive extracellular vesicles from a subset of endothelial progenitor cells rescue retinal ischemia and neurodegeneration. JCI Insight 7, (2022).

